# Epigenome Alterations and 3D Chromatin Architecture Remodeling in Inflammatory Macrophage Activation under Diabetic Conditions

**DOI:** 10.64898/2026.05.27.727961

**Authors:** Marpadga A Reddy, Supriyo Bhattacharya, Vinay Singh Tanwar, Vajir Malek, Zhuo Chen, Lingxiao Zhang, Linda Lanting, Xiwei Wu, Maria B. Grant, Joshua Tompkins, Zhen Bouman Chen, Rama Natarajan

**Affiliations:** Department of Diabetes Complications & Metabolism; Department of Diabetes & Cancer Discovery Science, Arthur Riggs Diabetes & Metabolism Research Institute and Beckman Research Institute of City of Hope, Duarte, CA; Department of Medical Oncology and Therapeutics Research, Beckman Research Institute of City of Hope, Duarte; Department of Opthalmology and Vision Science, University of Alabama at Birmingham, Birmingham, AL

**Keywords:** 3D chromatin architecture, Diabetes, Macrophages, Inflammation, Enhancer connectome, CRISPR-interference

## Abstract

Aberrant monocyte/macrophage activation in diabetes drives chronic inflammation and diabetic complications. Although epigenetic mechanisms are implicated in these processes, the contribution of three-dimensional (3D) chromatin reorganization remains unclear. Here, we use integrated multi-omics approaches to profile gene expression, chromatin accessibility and 3D chromatin architecture in human CD14⁺ monocyte-derived macrophages exposed to high glucose and TNF-α (HT), mimicking the diabetic milieu. HT induces inflammatory transcription programs that recapitulate those observed in diabetes and dynamically remodel chromatin accessibility, enhancer-promoter interactions, and topologically associated domains. Inflammatory genes exhibit increased chromatin accessibility, whereas cell cycle and metabolic genes display reduced accessibility and weakened enhancer-promoter interactions. These architectural changes promote the convergence of lineage-determining and signal-dependent transcription factors at enhancer-promoter loops, generating distinct regulatory hubs that orchestrate pro-inflammatory responses. Similar enhancer-promoter interactions are observed in peripheral blood derived monocytes from individuals with diabetes. Furthermore, perturbing chromatin interactions via CRISPR interference targeting HT-induced enhancers suppresses nearby *CCL2* and other chemokine genes. Collectively, this first comprehensive enhancer connectome map of macrophages under diabetic conditions advances the field by revealing extensive disease-associated rewiring of the 3D epigenome, highlighting chromatin architecture and epigenetic mechanisms as potential therapeutic target for diabetes-associated inflammation.

**Graphical abstract:** 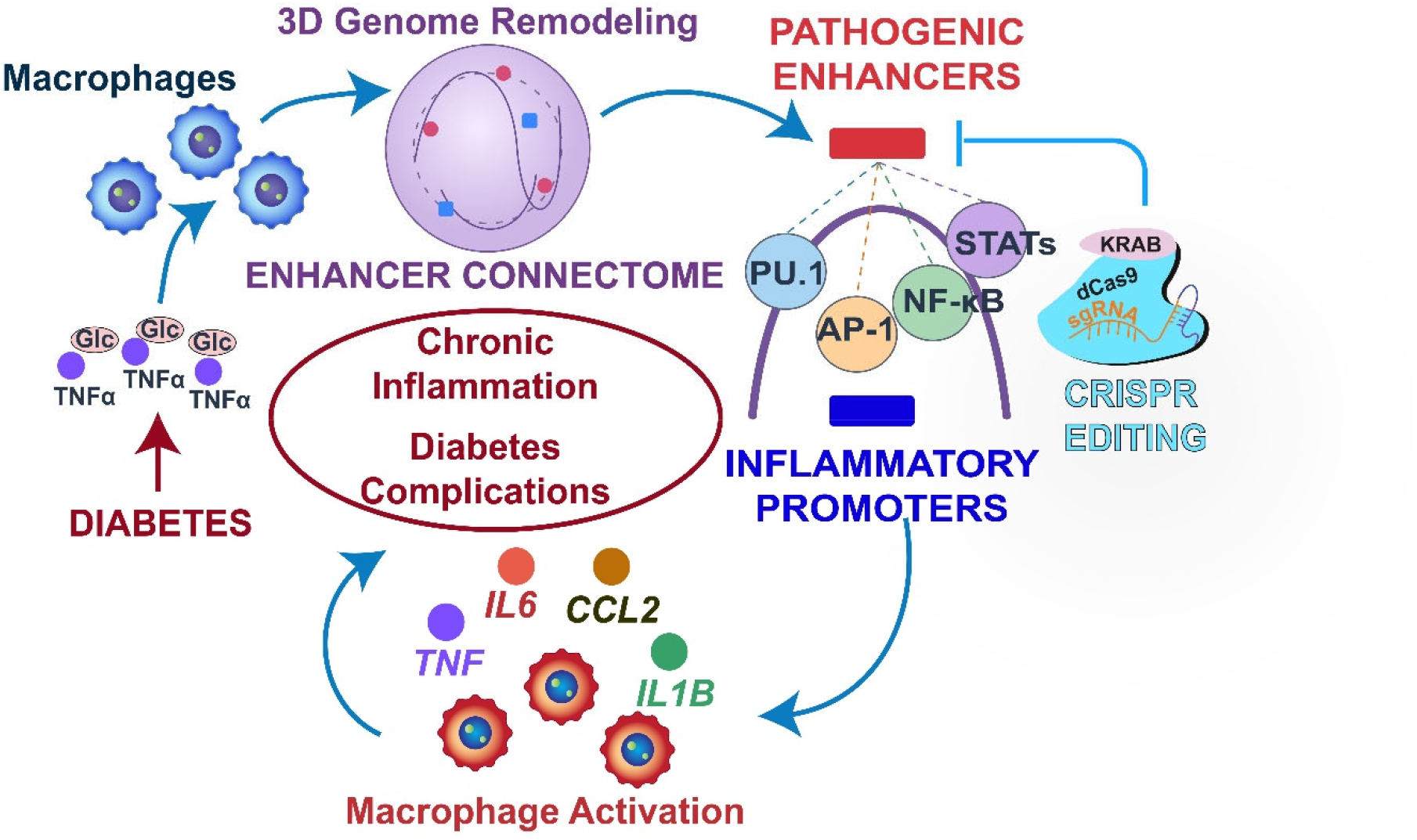

In diabetes, high glucose (Glc) and inflammatory cytokines remodel macrophage 3D chromatin architecture and enhancer connectomes, altering enhancer-promoter interactions, facilitating the convergence of lineage-specific and signal-dependent transcription factors (such as NF-κB) that regulate inflammatory factors associated with macrophage activation. This can promote chronic inflammation and accelerate diabetic complications. Targeted enhancer editing strategies could enable new epigenetic therapies to mitigate chronic inflammatory diabetes complications.

## 1. INTRODUCTION

Diabetes is associated with chronic inflammation, a key driver of the accelerated development of multiple macro- and microvascular complications such as atherosclerosis, hypertension, diabetic kidney disease (DKD), and diabetic retinopathy (DR)(*1*) . A key feature of diabetes is its impact on the innate immune system, particularly monocytes and macrophages, which play essential roles in inflammation, tissue repair, and pathogen defense (*2–5*). Chronic low-grade inflammation associated with diabetes results in the accumulation and activation of macrophages in metabolic tissues, contributing to pancreatic islet dysfunction, insulin resistance, hyperglycemia, and increased production of free fatty acids and inflammatory cytokines (*3–5*). Furthermore, accumulation and activation of macrophages in vascular, renal and other target tissues during diabetes augment the development of inflammatory vascular complications such as atherosclerosis and DKD (*1, 5, 6*).

Monocytes and macrophages are highly adaptable and can alter their phenotype by integrating diverse simultaneous signals to generate an appropriate inflammatory response (*2*). In diabetes, monocytes and macrophages are exposed to a complex pro-inflammatory milieu consisting of high glucose (HG), advanced glycation end products, inflammatory cytokines [i.e., tumor necrosis factor-alpha (TNF-α)], and free fatty acids, which skew macrophages towards pro-inflammatory M1-like and defective alternatively activated phenotypes (*7*). This disrupts inflammation resolution and tissue repair, which is further exacerbated by increased monocyte counts (monocytosis). In addition, dysregulation of macrophage metabolism, including a shift toward aerobic glycolysis (the Warburg effect) (*8*), and innate immune cell memory (called trained immunity) in response to hyperglycemia, oxidized lipids and insulin resistance(*9*) can also contribute to pro-inflammatory macrophage activation. The innate immune cell plasticity is driven by multiple factors, such as cell surface receptors recognizing chemokines (e.g. C-C Motif Chemokine Ligand 2 (CCL2) and CXCL10) and inflammatory cytokines (e.g. TNF-α and Inteleukin-1beta [IL-1β]), leading to activation of intracellular kinases and downstream transcription factors (TFs) like NF-κB) (*1*). Additionally, epigenetic mechanisms (e.g., DNA methylation, histone modifications, and non-coding RNA actions) have been implicated in sustained transcriptional responses to proinflammatory signals (*10*).

Within any given eukaryotic cell, gene transcription is modulated by TFs and histone-modifying and chromatin-remodeling proteins, which coordinate to regulate chromatin structure and the epigenetic landscape. These can lead to increased interactions between distant DNA regulatory elements known as enhancers and proximal gene promoters to activate gene expression, or transitions between heterochromatin (inactive, condensed, inaccessible to TFs) and euchromatin (active, open and accessible to TFs and associated with transcriptional activation) (*11–14*). Aforementioned epigenomic changes can now be interrogated using innovative chromatin conformation capture technologies that reveal higher-order chromatin organization of active open chromatin (A-type) and inactive closed chromatin (B-type) compartments (*15, 16*), composed of clusters of topologically associated domains (TADs) separated by boundaries (*17*). This complex 3D genome architecture enables fine-tuned and precise control of gene regulation expression by not only regulatory interactions among TFs at proximal promoters, but also establishing short- and long-range chromatin interactions and loops among enhancers and promoters in response to diverse stimuli (*17*). Enhancers mostly interact with promoters within the same TAD, and disruption of TAD boundaries leads to dysregulated gene expression in pathological conditions (*18, 19*). Chromatin interactions between enhancers and promoters can be rapidly formed, allowing rapid and efficient activation of TFs and gene expression during inflammatory responses (*20*). Dynamic alterations in these interactions and formation of *de novo* chromatin loops have been reported in response to microbial agents and inflammatory agents/cytokines, suggesting chromatin flexibility during immune responses (*21, 22*). During the inflammatory response, such as that induced by diabetic stress, macrophages undergo rapid and profound transcriptome-wide changes. However, little is known if and how changes in 3D genome architecture regulate monocyte/macrophage dysfunction associated with chronic inflammation in diabetes.

Here, we performed integrative multi-omics analyses to investigate the 3D chromatin remodeling and the related epigenetic mechanisms that drive proinflammatory macrophage activation in diabetes. We differentiated non-diabetic human peripheral blood CD14⁺ monocytes into macrophages and treated them with high glucose (HG) plus TNF-α (HT) to mimic the proinflammatory diabetic milieu. We then profiled the macrophage transcriptome using RNA-seq, chromatin accessibility using Assay for Transposase-Accessible Chromatin (ATAC-seq), and enhancer connectome using H3 lysine27-acetylation (H3K27ac, enhancer mark)-mediated chromatin immunoprecipitation (ChIP-seq) and HiChIP. HT treatment induced dynamic pro-inflammatory transcriptional programs, orchestrated by extensive changes in chromatin accessibility and reorganization of 3D chromatin, including alterations in enhancer-promoter interactions and disruption of TAD boundaries. We identified TF network hubs comprising interactions among lineage-specific enhancer-bound pioneer and signal-dependent TFs with promoter-proximal TFs at inflammatory gene loci, coincident with phenotypic changes associated with distinct macrophage responses. Similar HT-induced signaling pathways and enhancer-promoter interactions were observed in peripheral blood-derived monocytes from volunteers with type 2 diabetes (T2D) relative to healthy controls. CRISPR-interference targeting HT-induced enhancers suppressed expression of nearby chemokine genes like *CCL2, CCL1* and *CCL7* in macrophages, demonstrating the importance of these chromatin conformation changes. These findings reveal disease-associated rewiring of the macrophage 3D epigenome landscape and TF regulatory networks in diabetes that contribute to a sustained proinflammatory macrophage phenotype. They also highlight epigenetic mechanisms as potential therapeutic targets for inflammatory diabetic complications.

## 2 RESULTS

### 2.1 High glucose (HG) plus TNF-*α* (HT) induces extensive changes in the transcriptome of macrophages derived from human CD14^+^ monocytes

In diabetes, HG and inflammatory cytokines like TNF-α (TNF) play important roles in the adverse activation of monocytes and macrophages. Previous studies have shown that HG plus TNF-α (HT) treatment in part recapitulates key features of the diabetic inflammatory milieu in vitro in vascular and immune cells(*23–26*). Therefore, we treated human macrophages derived from CD14^+^ monocytes isolated from peripheral blood mononuclear cells (PBMCs) of healthy donors with HT (25 mM glucose +TNF) or NG (5.5 mM glucose) to examine changes in 3D chromatin architecture and epigenetic mechanisms associated with macrophage activation under diabetic conditions. We differentiated CD14^+^ monocytes derived from the peripheral blood mononuclear cells (PBMCs) of healthy donors into macrophages using macrophage colony-stimulating factor 1 (50 ng/ml, for 7 days). In an initial test, using human macrophages from a healthy donor, we found that HT significantly enhanced *IL1B* and *CCL2* expression compared with TNF alone i.e. NG+TNF (NT) treatment, establishing that HG amplifies the pro-inflammatory effects in these PBMC-derived macrophages (**Fig. S1A-B**) and supporting our rationale of using HT to mimic the effect of diabetes. We then treated macrophages from 3 non-diabetic donors with HT (**Fig. 1A**) and validated *IL1B* induction, a canonical inflammatory gene marker (**Fig. S1C**). To query the genome-wide regulatory mechanisms dictating gene expression in HT-macrophages, we performed integrative multiomics assays, including RNA-seq, ATAC-seq, and ChIP-seq and HiChIP with an H3K27ac antibody (**Fig. 1A** and **Methods**). For RNA-seq and ATAC-seq, samples from three separate donors were used. For H3K27ac-HiChIP, we used chromatin from one of the 3 donors, as well as pooled chromatin from the remaining two donors to obtain adequate input, yielding Sample 1 and Sample 2 for HiChIP respectively. For H3K27ac-ChIP-seq, we pooled remaining chromatin after HiChIP from all three samples. Collectively, these modalities generated a comprehensive map of the 3D chromatin architecture in macrophages and enabled us to evaluate the role of chromatin changes/rewiring in the dysregulated macrophage gene expression under the pro-inflammatory, diabetic milieu (**Fig. 1A**).

**Fig. 1.**
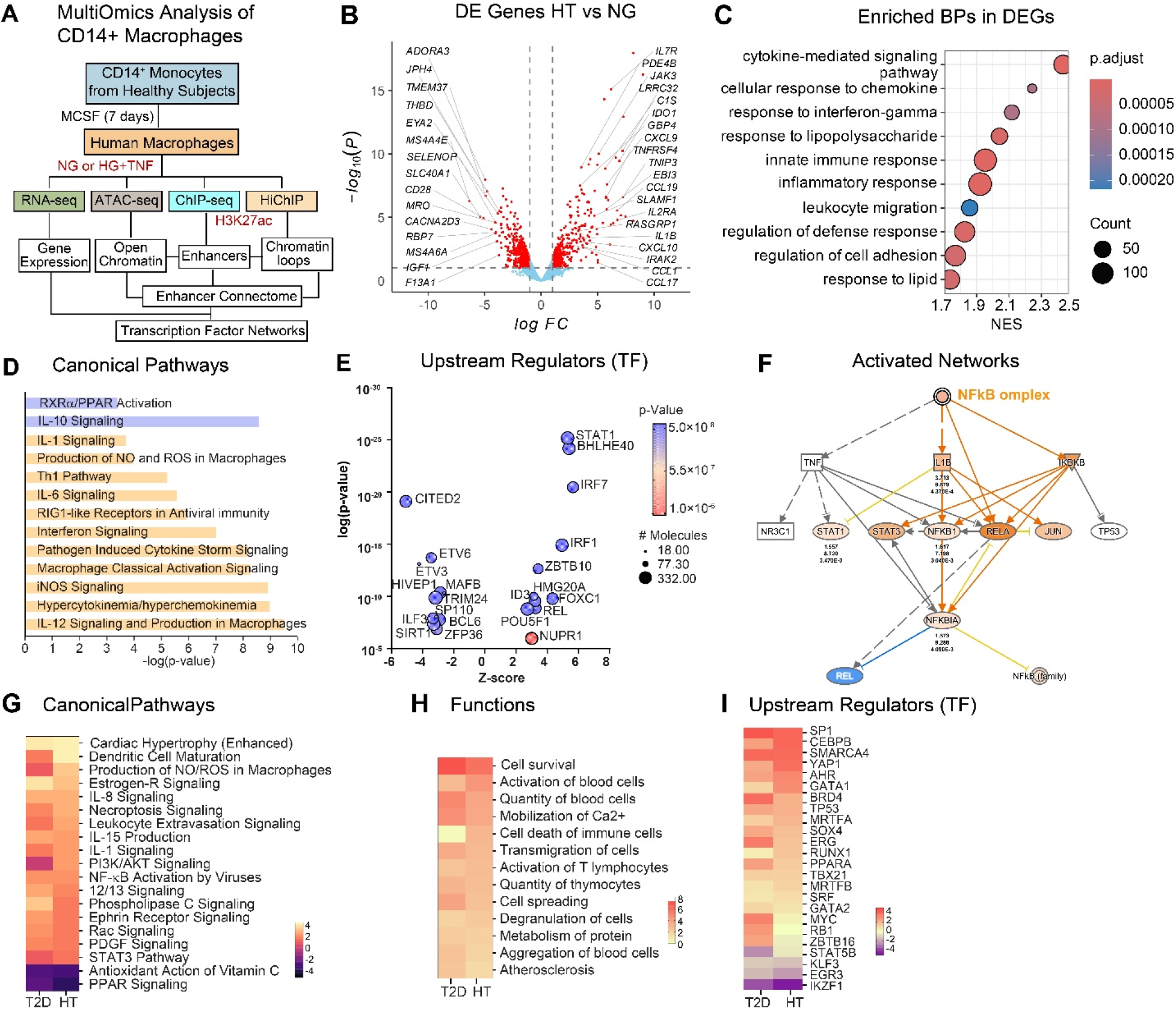
Activation of pro-inflammatory gene networks under diabetic conditions in human macrophages. **A.** Schematic of comprehensive multi-omics integrative analysis of human macrophages. CD14^+^ monocytes from healthy individuals were differentiated into macrophages with MCSF (50 ng/ml, 7 days), and treated with normal glucose (NG) or high glucose (HG) + TNF-α (HT) for 3 days. In all the indicated assays, results were compared between HT- and control NG-treated macrophages. **B.** Volcano plot of differentially expressed genes (DEGs) identified in RNA-seq in HT vs NG-treated human macrophages (n=3). **C.** Enriched biological processes (BPs) in DEGs from Gene Ontology-Biological Process (BP). **D-E.** Results of Ingenuity Pathway Analysis (IPA) showing canonical pathways (**D**, upregulated pathways in HT in orange, and downregulated ones in blue) and Upstream Regulatory Transcription factors (TF) enriched in genes upregulated by HT (**E**). **F**. Activation of NF-kB TF networks in HT vs NG-treated macrophages. **G-I.** Comparison analysis (using IPA) of canonical pathways (**G**), Functions (**H**) and Upstream regulatory TFs (**I**) enriched in DEGs between HT (vs. NG) treated human macrophages from this study versus CD14^+^ monocytes from T2D patients (vs healthy controls) RNA-seq data sets from a previous study (*25*).

RNA-seq showed that key M1 and M2 macrophage phenotype genes were increased and decreased respectively, in cells treated with HT compared with NG, indicating an enhanced inflammatory phenotype in these macrophages (**Fig. S1D**). Furthermore, Gene Set Enrichment Analysis (GSEA) of differentially expressed genes (DEGs) in HT vs NG (**Fig. 1B**) showed that upregulated genes were enriched with Gene Ontology (GO) biological processes (BP) associated with macrophage inflammatory response such as cytokine mediated signaling, chemokine response, and innate immune responses to interferon gamma, lipopolysaccharide (LPS) and lipids and (**Fig. 1C**). Consistently, Ingenuity Pathway Analysis (IPA) identified activation of canonical pathways that mediate pro-inflammatory signaling associated with cytokine storm, macrophage classical activation, and activation of signaling by iNOS, Interleukins and interferons. In contrast, anti-inflammatory or alternative activation signaling induced by IL-10 and RXRa/PPAR were inhibited (**Fig. 1D & S1E**). These changes were accompanied by the activation of pro-inflammatory transcription factors (TFs), including NF-κB/REL, interferon regulatory factors (IRFs), and signal transducers and activators of transcription (STATs), along with suppression of key anti-inflammatory and homeostatic regulators such as CITED2, SIRT1, BCL6, and CBX5 (HP1α), as well as TFs promoting alternative macrophage activation, including MAFB, (**Fig. 1E and Fig. S2A-B**). This network of TFs corresponded to the NF-κB complex activation, well established to be associated with increased expression of inflammatory cytokines, and collaboration with other TFs (**Fig. 1F**).

We then compared the HT-induced transcriptomic changes in macrophages with those detected in CD4^+^ monocytes from individuals with T2D (versus non-diabetic controls, HbA1c 9.32±2.4 vs 5.02±0.11, n=5 each), using a T2D dataset that we previously published (*27*). Analysis of the HT-upregulated genes revealed several shared pathways between the two datasets. These included activation of cytokines (IL-1, IL-8, and IL-15), nitric oxide/reactive oxygen species (NO/ROS), growth factor (PDGF), NF-κB, PI3K/Akt, and STAT3 signaling pathways and dendritic cell maturation, but suppression of PPAR and anti-oxidant signaling (*28*) (**Fig. 1G**). HT and T2D conditions also affected similar functions like cell survival and transmigration, calcium mobilization, immune cell activation, and blood cell aggregation (**Fig. 1H**), many of which are associated with atherosclerosis (*29*). In addition, TFs that play essential roles in monocyte/macrophage lineage, differentiation, and inflammatory phenotype, such as SP1, BRD4, TP53, RUNX1, MRTFA, KLF3, and EGR3 were also shared between the two datasets (**Fig. 1I**). The HT-downregulated genes showed enriched pathways related to hematopoietic progenitor development, cellular homeostasis, and fatty acid metabolism, myeloid cell adhesion, apoptosis, and others (**Fig. S3A-C).** As in the case of upregulated genes, comparison between HT treated and T2D patient cells revealed several shared enriched pathways and functions between downregulated genes in HT and T2D cells were associated with inflammatory, immune and insulin receptor signaling (**Fig. S3C**). Furthermore, Upstream Regulator analysis showed enrichment of several TFs like SMAD2, MAFB, CEBP family that regulate macrophage differentiation and alternative activation (**Fig. S3D**). Thus, HT treatment recapitulates key features of dysregulated monocytes and macrophages in human diabetes by activating multiple disease-relevant pathways and cellular functions.

### 2.2 ATAC-seq reveals HT induces extensive changes in chromatin accessibility in human macrophages

We next performed ATAC-seq to map open chromatin regions which are accessible to regulatory factors like TFs in response to HT. Analysis of differential ATAC-seq peaks (see Methods) identified 1279 regions with increased chromatin accessibility (upregulated peaks) and 1761 regions with reduced chromatin accessibility (downregulated peaks) in HT versus NG-treated macrophages. These were distributed among promoters, transcription start sites (TSS), gene body, and intergenic regions (**Fig. 2A**). HT-upregulated peaks corresponded to significant enrichment of immune and inflammatory responses, cytokine production, apoptotic and pattern recognition signaling pathways (**Fig. 2B**), in line with the inflammatory activation at the transcriptomic level. On the other hand, peaks with reduced chromatin accessibility (HT-downregulated peaks) were associated with regulation of hemopoiesis, leukocyte differentiation, JAK-STAT cascade, regulation of p38MAPK cascade, and circadian rhythm (**Fig. S4A**). Integrated analysis of the RNA-seq and ATAC-seq data revealed that genes involved in these pathways indeed possess differential chromatin states in HT-treated cells (**Fig. 2C**). For example, HT treatment increased chromatin accessibility near the upregulated pro-inflammatory gene *IL1B* (**Fig. 2D**). In contrast, chromatin accessibility decreased in HT near downregulated *MAFB,* which regulates alternative activation (M2-like phenotype) of macrophages (**Fig. 2D**). Furthermore, ATAC-seq peaks in our datasets matched those of open chromatin regions identified by DNase-seq in ENCODE data, corroborating credence to our observations (**Fig. 2D**).

**Fig. 2.**
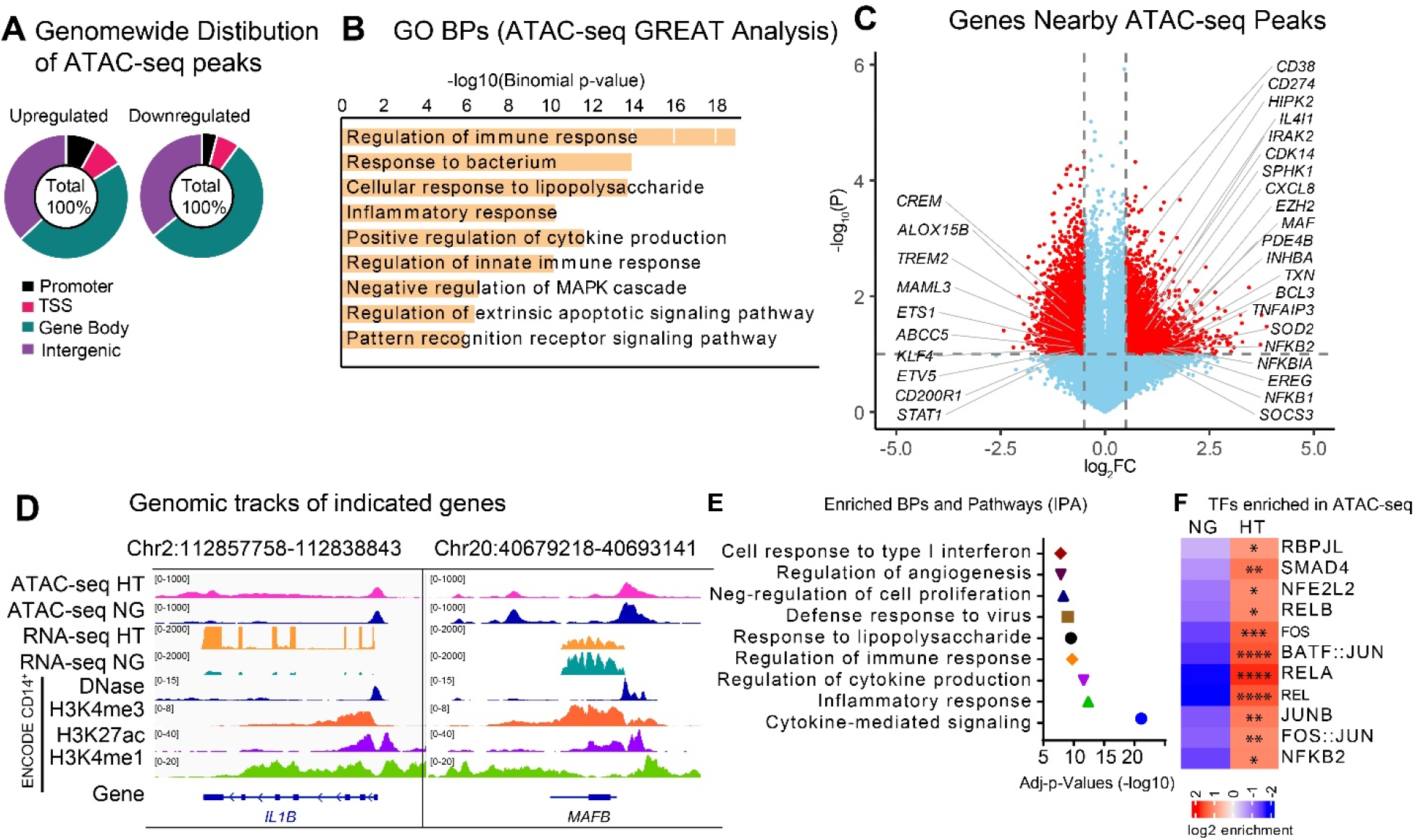
ATAC-seq reveals extensive changes in chromatin accessibility induced by HT in human macrophages. **A.** Genomic distribution of open chromatin (ATAC-seq data) in human macrophages treated with HT and NG (n=3). Human CD14^+^ monocytes were differentiated into macrophages, treated with NG or HT, and subjected to ATAC-seq analysis. **B.** Results of GREAT analysis showing Gene Ontology Biological Processes (GO BPs) enriched in ATAC-seq peaks. **C.** Integrated analysis of RNA-seq and ATAC-seq data showing differentially regulated genes near open chromatin. **D.** Integrated genomic view (IGV) of ATAC-seq and RNA-seq tracks of upregulated gene *IL1B* and downregulated gene *MAFB* in HT vs NG. For ease of visualization, RNA-seq and ATAC-seq data tracks from 3 samples each (NG and HT) have been combined using IGV tools. Also shown are DNase-seq, and ChIP-seq (H3K4me3, H3K27ac and H3K4me1) in human CD14^+^ monocytes from ENCODE datasets. **E.** GO BPs (BPs) enriched in differentially expressed genes near ATAC-seq peaks. **F.** Transcription factor (TF) motifs enriched in differentially regulated (Upregulated) open chromatin. Only TF-motifs that are shared between RNA-seq and HiChIP are shown. p.adj: *p<0.1-0.01; **p<0.01-0.001; ***p<0.001-0.0001.

DEGs near regions with HT-increased chromatin accessibility revealed enrichment of pathways related to signaling by interferons, viruses, regulation of cytokine production as well as inflammatory response (**Fig. 2E**). Supporting this, promoter regions with increased accessibility were enriched in TF motifs for NF-κB (RELA, RELB and NF-kB2) and AP1 (Fos and Jun), and BATF:JUN, and interferon regulatory factors (IRFs), (**Fig. 2F),** whereas TF motifs like ETV2, HIF1A and MAFA were enriched in downregulated peaks (**Fig. S4B**), key TFs associated with macrophage functions involved in immune responses and inflammation suppression (*30–34*). Overall, the integrated analysis of ATAC-seq and RNA-seq indicated a coordinated modulation of chromatin accessibility and TF activity at inflammatory loci in HT-treated macrophages, which can contribute to HT-regulated inflammatory gene expression and chronic inflammation in diabetes.

### 2.3 HiChIP analysis reveals disruption of 3D genome organization by HT in human macrophages

The profound genome-wide changes in accessible regulatory elements induced by HT prompted us to further investigate diabetes-induced rewiring of the 3D chromatin architecture in human macrophages. We analyzed HiChIP and ChIP-seq targeting H3K27ac to assess enhancer-promoter loops and large-scale changes in chromatin architectural features in HT, including topologically associated domains (TADs). For HiChIP, Samples 1 and 2 were processed and analyzed together. Because Sample 1 exhibited superior peak resolution and a higher signal-to-background ratio compared with Sample 2, data from Sample 1 were used for primary visualization and analysis of HT-induced changes. The principal findings, such as TAD boundary disruptions and direction of loop changes at key gene promoters were subsequently confirmed and validated using Sample 2 as an independent biological replicate.

The average number of TADs per chromosome was higher (67 vs 51, p=0.037) and TAD lengths were shorter (1.8 vs 2.5 Mb, p=0.002) in HT compared with NG treated macrophages (**Fig. 3A-B**), suggesting genome-wide TAD disruption by HT. Integration with RNA-seq data linked these structural changes to altered expression of genes associated with inflammatory macrophage phenotype. This was also underscored by the changes in gene expression at TAD boundaries on multiple chromosomes, and many of these genes regulate inflammatory responses (**Fig. 3C**). Notably, *NFKBIA*, a key gene involved in inflammatory and TNF signaling, is located at one of these disrupted TADs on chromosome 14 (**Fig. 3D**). Nearby located genes *RTN1* and *PSMA6*, which are involved in ER stress (*32*), were also upregulated by HT in macrophages. Pathway analysis using genomic regions with HT-disrupted TAD boundaries showed enrichment of inflammatory signaling pathways (**Fig. 3E**) and TF motifs (**Fig. 3F**). Furthermore, we validated these HT-induced changes in TAD disruption using the replicate sample (sample 2), in which HiChIP was performed using pooled chromatin from two donors. This replicate data confirmed similar decreased TAD sizes, increased TAD numbers (**Fig. S5A-B**), TAD disruption at *NFKBIA* locus on Chr14 (**Fig. S5C**), and enrichment of inflammatory GO BPs at disrupted TADs (**Fig. S5D**). Moreover, we identified 112 HT-disrupted TAD boundaries shared between the donor samples 1 and 2 (**Fig. S5E**) that were enriched for key inflammatory pathways including IL2-STAT5 and TNFα signaling, hypoxia, and inflammatory response (**Fig. S5F**). We also used publicly available paired HiC and CTCF ChIP-seq datasets from human macrophages differentiated using MCSF (*35*) to confirm our HiChIP-derived TAD boundaries at the *NFKBIA* locus, which showed similar CTCF enrichment at the *NFKBIA* locus in NG-treated macrophages versus M0 macrophages (**Fig. S6A**). In addition, the genome-wide distribution of TADs in sample 1 and sample 2 showed significant overlap with each other and with M0 macrophages in this dataset, further validating our TAD analysis (**Fig. S6B**). These changes in TADs are consistent with inferences drawn from the RNA-seq, ATAC-seq and ChIP-seq data, indicating significant remodeling of the 3D chromatin architecture under HT mapped by our pioneering high-resolution HiChIP data, and constitute a critical epigenetic mechanism contributing to chronic inflammation in diabetes.

**Fig. 3.**
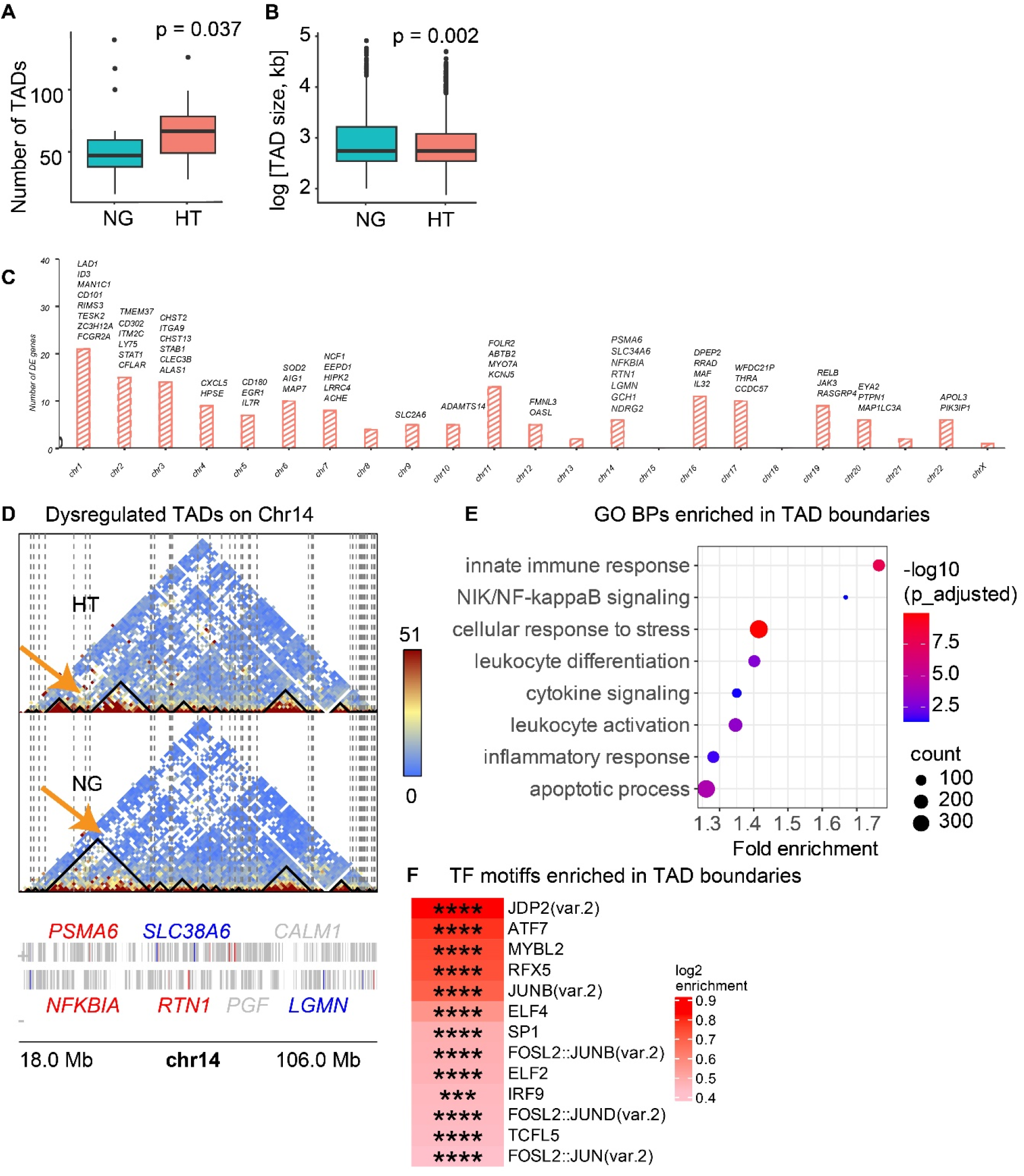
Disruption of hierarchical 3D genome organization by HT in human macrophages. **A-B.** HT shortened the TAD lengths (**A**) and increased the number of TADs per chromosome (**B**) vs NG. This data represents HiChIP analysis results of Sample 1 (single donor). **C**. TAD-associated differentially expressed (DE) genes per chromosome. **D.** Heatmap of normalized HiChIP read density in HT vs NG on chr 14. TADs are highlighted by black triangles and disrupted TAD boundaries by dashed vertical lines. HT-regulated genes are highlighted in red (up) and blue (down) colors. **E**. Pathways regulated by genes in TAD boundaries disrupted by HT in human macrophages. **F.** TF motifs enriched at disrupted TAD boundaries. p.adjusted: ***p<0.001-0.0001; ****p< 0.0001.

### 2.4 HT induces extensive changes in the enhancer connectome in human macrophages

The HT-induced changes in TAD boundaries indicate genome-wide alterations in chromosomal interactions. A key mechanism for gene regulation involves interactions between enhancers and promoters, or the enhancer connectome (*14, 36*). To this end, we used integrated H3K27ac HiChIP and ChIP-seq datato map active enhancers and their promoter interactions, enabling the characterization of dynamic chromatin looping and the construction of an enhancer connectome (*36*) in NG- and HT-treated macrophages. HiChIP data was analyzed using the HiCDC^+^ pipeline (*37*) to identify chromatin loops and differential regulation in HT versus NG at 5kb resolution (see Methods). This effort generated the first comprehensive Enhancer Connectome and *in situ* HiChIP maps of DNA loops altered under diabetic conditions in human macrophages.

We identified a total of 1,275,075 chromatin looping events in HT-treated and 876,842 loops in NG-treated macrophages, with 266,510 loops (qvalue < 0.05) shared between the two groups (**Fig. S7A**). The identified loops were mostly aligned to enhancers (42%) and promoters (21%), with the remaining aligned to gene body and intergenic regions (**Fig. S7B**). Comparing HT versus NG, 24,233 differentially regulated chromatin loops (logFC > 1) were identified.

Integrative analysis of Enhancer connectome with ATAC-seq and RNA-seq datasets revealed a significant genome-wide correlation between differentially regulated enhancer-promoter loops and open chromatin with changes in gene expression (**Fig. 4A-B**). The highest expression changes in HT were observed in genes associated with both altered chromatin accessibility and enhancer loops (**Fig. 4A)**, exemplified by upregulated genes like *TNIP1* and *CD38* that were associated with increased enhancer-promoter loops and open chromatin. In contrast, downregulated genes (e.g., *ADAM19*, *CREM*) were associated with reduced chromatin accessibility and enhancer-promoter interactions (**Fig. 4B**), underscoring a critical role for enhancer-mediated looping in HT induced gene regulation. Genomic visualization of HiChIP tracks at upregulated inflammatory gene loci such as *IL1B* (**Fig. 4C)***, NFKBIA* and *TNIP1* (**Fig. S8 A-B**) showed increased enhancer loops in HT versus NG, accompanied by increased open chromatin, whereas downregulated antioxidant and immune suppressor genes such as *SELENOP* and *CD300A* showed decreased enhancer loops in HT (**Fig. S8 C-D)**. In further support of this, genes whose promoter regions were involved in enhancer loops in HT showed enrichment of inflammatory and immune response processes (Genomic Regions Enrichment of Annotations Tool [GREAT] analysis; **Fig. S7C**). In contrast, downregulated genes involved in metabolism, cell cycle, IL-10 production and macrophage apoptosis were associated with decreased chromatin accessibility and enhancer looping (**Fig. S7D**). Furthermore, enhancer regions involved in HT-increased chromatin loops were enriched in binding motifs of TFs NF-kB, RELA and AP-1 (FOS/JUN) (**Fig. 4D**), while HT-decreased loops were associated with cell cycle and stress response TFs such as ATF1, ATF7 and MTF1 (**Fig. S7E**), in agreement with the ATAC-seq and RNA-seq data.

**Fig. 4.**
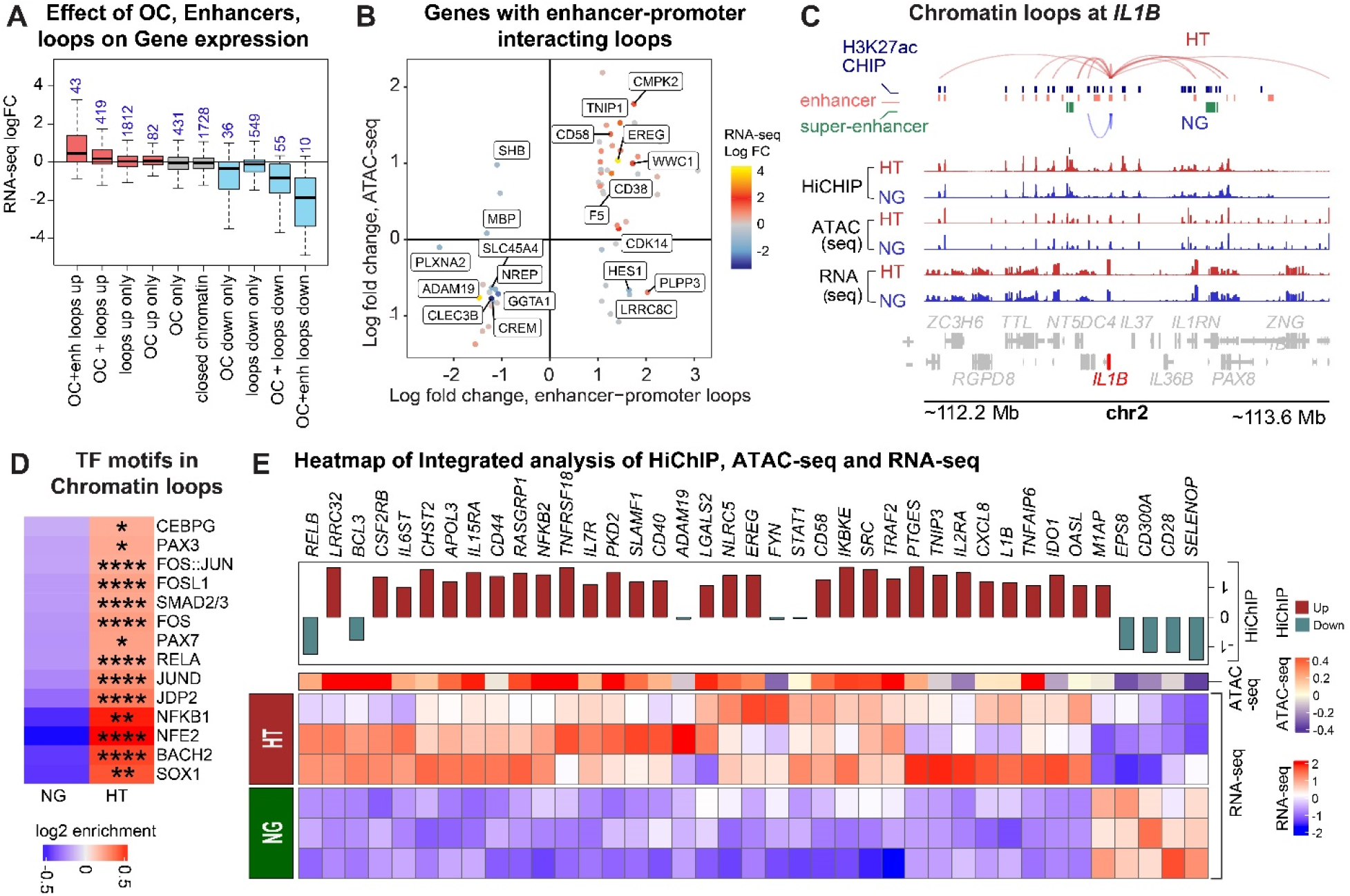
HT induced dynamic changes in the enhancer connectome in human macrophages. **A.** Results of HiChIP analysis showing the effect of Open chromatin (OC), enhancers, and chromatin loops on gene expression. HiChIP with an H3K27ac antibody was performed in human CD14^+^ monocytes (from a single donor) differentiated into macrophages and treated with NG or HT, and the results were overlaid with chromatin accessibility (ATAC-seq) and gene expression (RNA-seq) data sets. **B**. Correlation between open chromatin and chromatin loops interacting with enhancers and promoters on gene regulation. **C.** Genomic view of HiChIP tracks showing enhancer-promoter chromatin loops at inflammatory gene *IL1B* in HT-treated (maroon color loops) versus NG-treated (blue color loops) human macrophages. The genomic tracks of open chromatin (ATAC-seq peaks) and gene expression (RNA-seq data) have also been shown. **D**. TF motifs enriched in HiChIP loops in HT vs NG treated macrophages. Similar TF motifs are also enriched in RNA-seq and ATAC-seq datasets. p.adj: *p<0.1-0.01; **p<0.01-0.001; ***p<0.001-0.0001; ****p< 0.0001 (hypergeometric test). **E.** Integrated analysis of HiChIP, ATAC-seq and RNA-seq. Heatmaps comparing the average normalized expression of key inflammation genes in HT vs NG, along with the logFC in ATAC-seq (OC) and HiChIP read densities in chromatin loops at indicated genes. Each row is scaled separately using the logFC of individual replicates. Results show a significant correlation between gene expression, open chromatin and chromatin loops interacting with enhancers and promoters.

The regulatory landscape that emerges upon integration of HiChIP, ATAC-seq and RNA-seq is summarized in **Fig. 4E**, highlighting the associative relationship between chromatin accessibility and enhancer looping with the induction of key inflammatory genes in HT (e.g*., IL1B, TRAF2, NFKB2*). As shown in the CIRCOS plot (**Fig. S9**), enhancer-promoter interactions involving inflammatory genes span multiple chromosomes, indicating genome-wide chromatin remodeling in HT. Moreover, we confirmed significant overlap in HT regulated HiChIP loops between sample 1 and the replicate sample 2, in which HiChIP was performed using pooled DNA from two donors as mentioned earlier (**Fig. S10 A-C**). In addition, sample 2 also showed similar changes in enhancer-promoter loops at key candidate loci upstream of *CCL2* (including ChemDE) and *IRAK2* (**Fig. S10D-F**). Collectively, HiChIP analysis and multiomics integration indicated that genome-wide enhancer-mediated chromatin loops mechanistically drive the pro-inflammatory gene program in HT-treated macrophages. The majority of DEGs and TFs involved in these chromatin conformation changes are known to play important roles in diabetic complications like DKD, DR and cardiovascular disease (CVD).

### 2.5 HT-induced promoter-enhancer looping at key inflammatory loci and a novel chemokine distal enhancer (ChemDE)

Overlaying publicly available enhancer information from multiple human cell lines in GenHancer SEdB datasets (*38*) along with our HiChIP data revealed increased HT-induced loop activity at several genomic loci with high density of known enhancers. Notably, we identified HT-induced chromatin loops linking distant enhancers to the cytokine *IL1B* (**Fig. 4C**), the chemokines *CCL2* and *CCL1*, and the key inflammation regulator IL-1R-associated kinase 2 (*IRAK2)*, all of which encode key pro-inflammatory molecules (**Fig. 5A-B)**. Among these, we identified a novel enhancer located ∼50 kilobases (kb) upstream of *CCL2* (chr17:34,200,277-34,203,135, blue-shaded, **Fig. 5A**), that exhibited significantly increased looping activity in HT. Based on GenHancer annotations, we noted that this unique enhancer is specific to macrophages and designated it as the <u>Ch</u>emokine <u>d</u>istal <u>e</u>nhancer (ChemDE). We observed that interactions of ChemDE with upstream regions of multiple chemokine genes (including *CCL2*) are enhanced in HT-treated human macrophages versus NG (**Fig. 5A**). Similarly, we found enhanced interactions between distal enhancers and the *IRAK2* promoter in HT versus NG (**Fig. 5B**). The HT-activated ChemDE was validated using PCR-based H3K27ac-ChIP and 3C chromatin conformation assays, along with HT-induced *CCL2* expression (**Fig. 5C-E**). The 3C assays also showed interactions between ChemDE and the *CCL2* upstream region in monocytes from T2D and non-diabetic individuals, although the difference was not statistically significant (**Fig. 5F).** Similar observations were made for *IRAK2* and the enhancer (chr3:10,164,750-10,203,595) that interacts with the *IRAK2* promoter (**Fig. S11A**), and notably 3C assays showed significant increase in these *IRAK2* genomic interactions in not only HT treated cells, but also T2D patient monocytes relative to control (**Fig. 5G-J**). In addition, 3C assays confirmed the HT-induced increases in chromatin-enhancer interactions at the *IL1B* and *NDRG1* loci (**Fig. S11B-C**) and in T2D samples (**Fig. S11D-E**). Collectively, these results suggest that HT induced genome-wide changes in the enhancer connectome and related epigenetic mechanisms can drive inflammatory gene activation in monocytes and macrophages under diabetic conditions.

**Fig 5.**
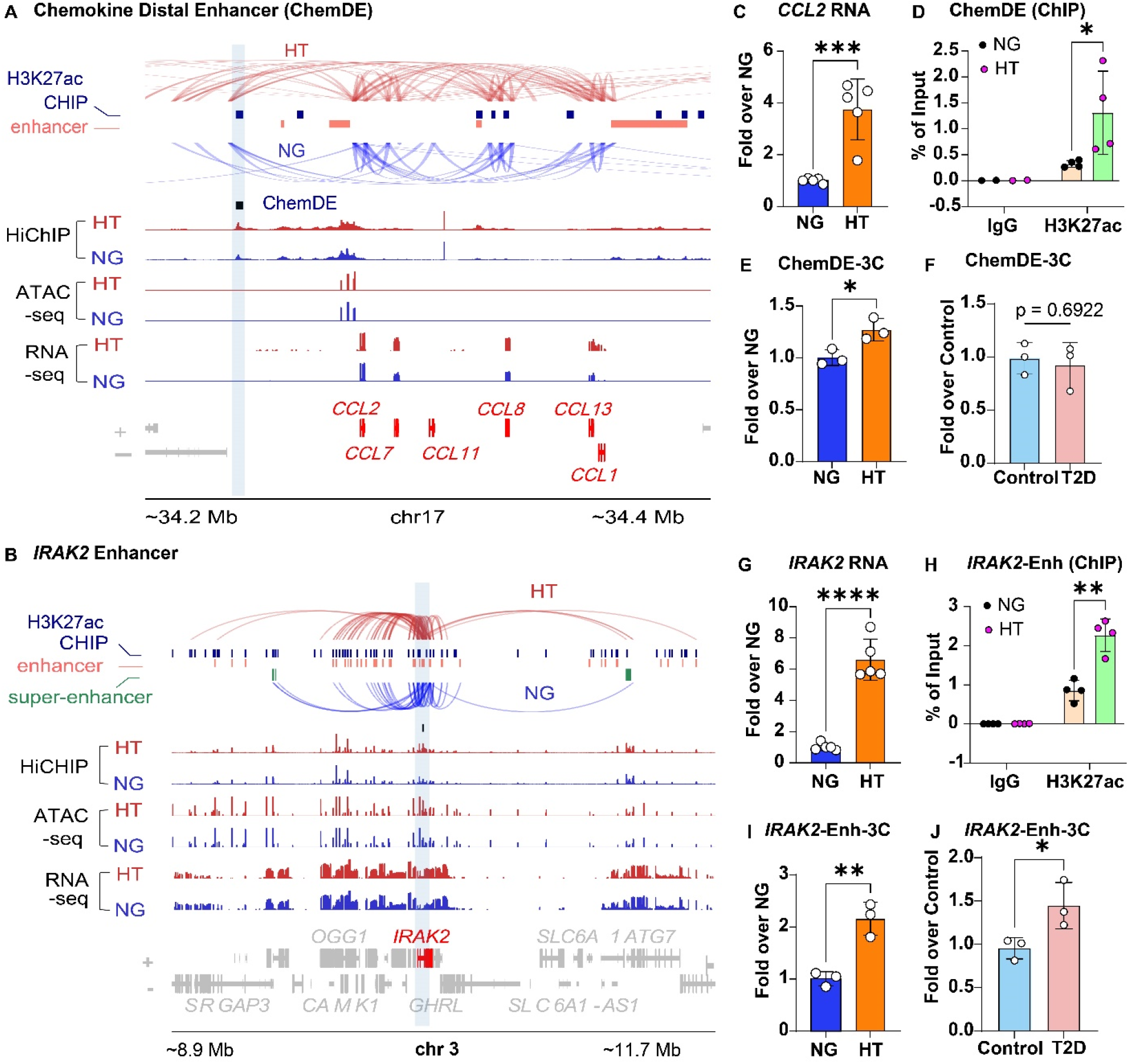
Identification of novel enhancers and chromatin loops regulated by HT and diabetes in human macrophages and monocytes. **A-B:** Genomic maps showing the interaction of chromatin loops between Chemokine distal enhancers (ChemDE) and promoters of chemokine genes on Chromosome 17 and *IRAK2* gene on chromosome 3. H3K27ac ChIP-seq, ATAC-seq and RNA-seq data tracks have also been shown. RNA-seq and ATAC-seq data tracks represent average of 3 samples each (NG and HT). **C-E.** Results of increased expression of *CCL2* by RT-qPCR (C), ChIP-qPCR analysis showing increased H3K27ac enrichment at ChemDE enhancer (D), and results of 3C assays showing ChemDE-*CCL2* promoter interactions in HT and NG treated macrophages. **F**. Results of 3C assay of ChemDE-CCL2 promoter interactions in monocytes from Control and diabetic (T2D) individuals. **G-I:** *IRAK2* expression **(G**), H3K27ac enrichment at the *IRAK2* enhancer (H), and 3C assays showing enhancer-promoter interactions (**I** and **S11A**) in HT treated macrophages. **J.** Results of 3C assays showing *IRAK2* enhancer-promoter interactions in cells from Control and T2D individuals. *, p<0.05, **, p<0.01, ***, p<0.001, ****, p<0.0001, as determined by unpaired t-test ( C, E-G, and I-J, n=3-5), and Multiple unpaired t-tests (D and H, n=4).

### 2.6 Regulation of HT-induced chemokine gene expression by ChemDE enhancer

Next, we investigated the functional roles of novel enhancers identified in this study, such as ChemDE in regulating HT-induced gene expression in human macrophages using CRISPR interference (CRISPRi) to target HT-induced enhancers. We used dCas9:SALL1:SDS3, a catalytically inactive dCas9 fused to the transcription repressive domains of Sal-like protein 1 (SALL1) and Sin3a corepressor complex component (SDS3) for CRISPRi. Targeting dCas9:SALL1:SDS3 to specific genomic locations with synthetic guide RNAs (sgRNAs) recruits repressive histone deacetylases and the Swi-independent three (SIN3) complex, leading to gene repression (**Fig. 6A**) (*39, 40*). We first optimized CRISPRi in THP1 monocytic cells by co-transfecting *in vitro* transcribed dCas9:SLL1:SDS3 mRNA along with sgRNAs targeting *PPIB* promoter (sgPPIB) and a non-targeting guide RNA (sgNTC) as control. Results showed inhibition of *PPIB,* but not *PPIA* expression by sgPPIB relative to sgNTC-transfected cells in monocytes and macrophages (**Fig. 6B-C**), Next, we targeted the novel enhancer ChemDE (chr17:34200277-34203135, Hg38), which interacts with promoter upstream regions of *CCL2* and other nearby chemokines (**Fig. 6D**). We designed sgRNAs targeting ChemDE (sgCDE) and co-transfected them with dCas9:SALL1:SDS3 mRNA in THP1 cells. Compared to sgNTC, sgCDE significantly attenuated HT-induced *CCL2* and *CCL1* (**Fig. 6E-F**), without affecting *CCL7* and *CCL11*, which are encoded in the gene cluster but not induced by HT in THP1 cells, or the control *PPIB* gene (**Fig. 6G-I**). Importantly, even in primary human macrophages derived from CD14^+^ monocytes of healthy donors, we confirmed similar inhibition of HT-induced chemokines *CCL2*, *CCL1* and *CCL7,* but not *CCL11* (which was not induced by HT) or control *PPIB*, by sgRNAs targeting ChemDE **Fig. 6J-N**). These results suggest that distantly located novel ChemDE interacts with chemokine gene promoters to regulate their expression in human macrophages under diabetic conditions.

**Fig. 6.**
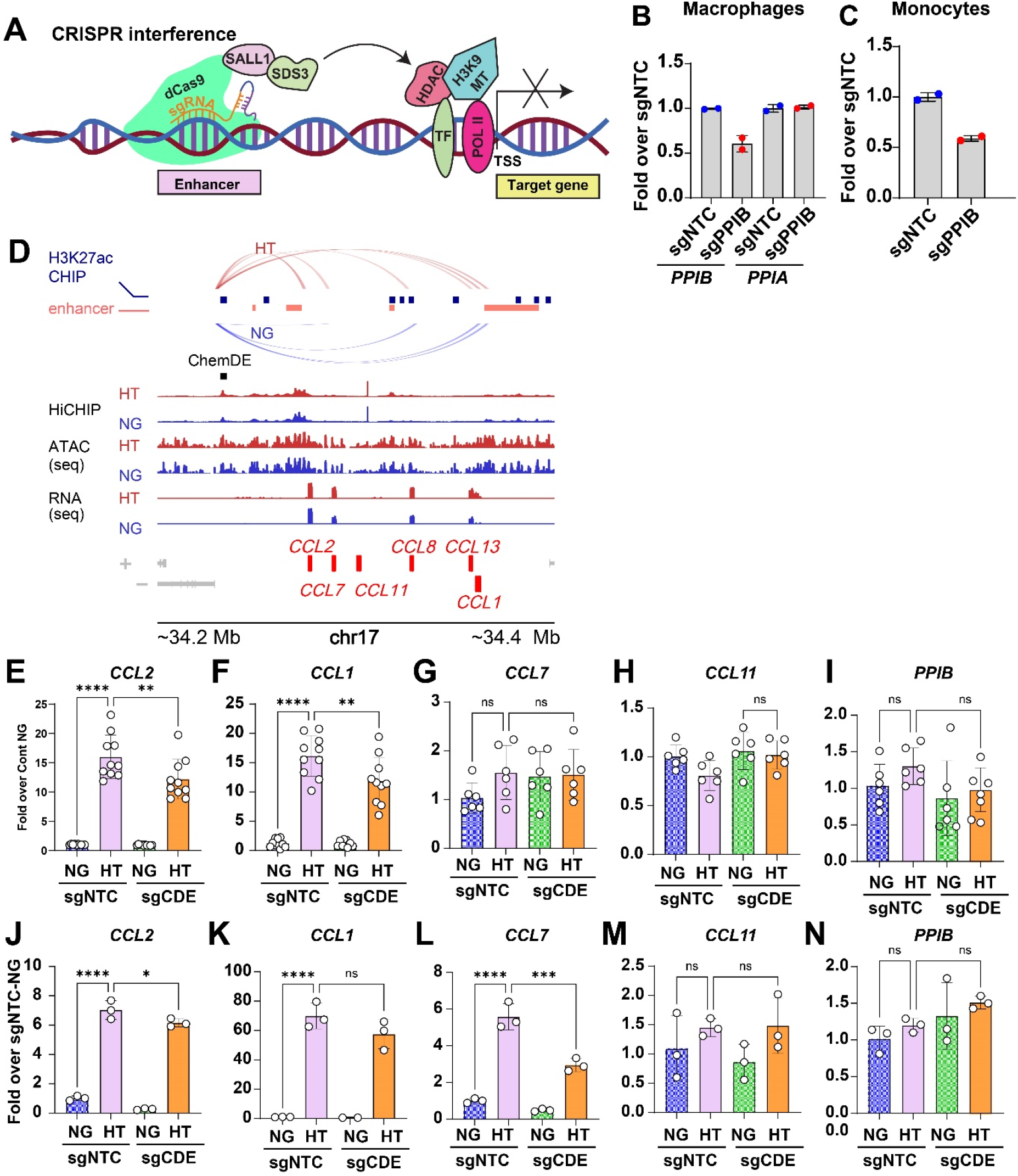
CRISPR interference (CRISPRi) demonstrates the function of Chemokine distal enhancer (ChemDE) in HT-induced chemokine gene regulation in human macrophages. **A.** CRISPRi using dCas9-SALL1 for epigenetic editing. Schematic of gene repression by targeting the indicated enhancers with catalytically inactive dCas9:SALL1:SDS3 using specific guide RNAs (sgRNAs), which recruit repressive histone-modifying enzymes to the target site, thereby inhibiting gene expression. **B-C.** Expression of indicated genes by RT-qPCR in THP1 macrophages and monocytes co-transfected with dCas9:SALL1:SDS3 mRNA along with *PPIB*-specific sgRNAs (PPIB) and non-targeting control sgRNA (sgNTC). **D.** Genomic views of ChemDE enhancer, HiChIP, ATAC-seq peaks, and RNA-seq of indicated chemokine loci in NG and HT treated human macrophages. Only the promoter interacting loops by the ChemDE enhancer are shown. Other enhancer-promoter loops in this region are depicted in Fig. 5A. **E-I.** Gene expression analysis (RT-qPCR) of indicated genes in THP1 macrophages co-transfected with dCas9:SALL1:SDS3 mRNA and sgNTC or ChemDE specific sgRNA (sgCDE) and treated with NG or HT. **J-N.** Gene expression analysis (RT-qPCR) of indicated genes in primary human macrophages co-transfected with dCas9:SALL1:SDS3 mRNA and sgNTC or ChemDE specific sgRNA (sgCDE) and treated with NG or HT. *, p<0.05, **, p<0.01, ***, p<0.001, ****, p<0.0001 as determined by Ordinary one-way ANOVA and Šídák’s multiple comparisons test (E-F, n=10; G-I, n=6; and J-N, n=3).

### 2.7 Identification of inferred TF networks driving HT-induced gene expression through enhancer-promoter interactions

Having demonstrated the central role of the enhancer connectome and 3D chromatin architecture rewiring in HT-induced macrophage gene expression/activation, we sought to determine the TF networks that are modulated by these changes (for details, see Methods). Using motif enrichment analysis, we identified TF motifs that were uniquely enriched in the promoter regions of HT-induced genes (designated as Promoter-motif), while others were only present in the enhancer regions with increased promoter loops in HT (designated as Enhancer-loop motif), suggesting distinct regulatory roles of these TFs in conjunction with the enhancer connectome (**Fig. S12A and Supplementary Tables S1-S2**). In addition, we used the upstream regulators analysis in IPA to predict the TFs regulating the differential gene expressions in HT (shown in **Fig. 1E**) and linked their motifs to the active promoter and enhancer regions. Comparison of IPA predicted TFs with enriched TF motifs in promoter and loop-mediating enhancer regions revealed a shared set of TFs enriched by HT at enhancer-loops, promoters and open chromatin regions, including upstream regulatory TFs identified by IPA (**Fig. S12B**). We henceforth focused on the most confident set of TFs involved in transcriptomic regulation in HT by combining the enhancer-loop associated TFs with the IPA predicted ones. Overlaying them onto the TF-target gene network using known protein interactions from the STRING database (*41*) revealed a complex and dense interactome involving both enhancer-bound and regulatory (promoter-bound) TFs **(Fig. S12C**). Notably, the enhancer bound TFs were enriched in pioneer factors capable of binding to condensed nucleosomes and opening the chromatin and recruiting other regulatory TFs to initiate transcription (*40*).

For an in-depth understanding of the TF network architecture in HT-macrophages, we clustered it into densely interconnected communities, revealing three distinct subnetworks involving key TFs that regulate distinct biological processes related to monocyte/macrophage differentiation, proliferation and inflammatory phenotype (**Fig. 7A-F**). Subnetwork 1 regulates cytokine signaling/secretion, innate immune responses and leukocyte proliferation, which can affect inflammatory phenotype and monocytosis in diabetes (**Fig. 7A and B**). Subnetwork 2 involves leukocyte migration, wound healing and apoptotic processes, hallmarks of activated macrophages and associated with inflammation (**Fig. 7C and D**). Conversely, Subnetwork 3 regulates key processes associated with defense responses and oxidative stress, which are crucial for host defense against pathogens, as well as extracellular matrix organization involved in adhesion and signaling by cytokines and growth factors, which are essential for macrophage functions (**Fig. 7E and F**). Interestingly, this analysis revealed interactions between enhancer bound pioneer TFs (green color, like SPI1 (PU.1), CEBP, MAF, SOX, FOXO), and signal dependent TFs (pink color, such as NF-κB, AP-1 (FOS/JUN), STAT, NFE2L2 (NRF2), BACH, ETS, IRFs) with promoter bound TFs (blue color) in all three Subnetworks (**Fig. 7A, C and E**). These results suggest importance of chromatin reorganization and looping in promoting interactions between TFs binding at distant enhancers with target promoters to regulate gene expression.

**Fig 7.**
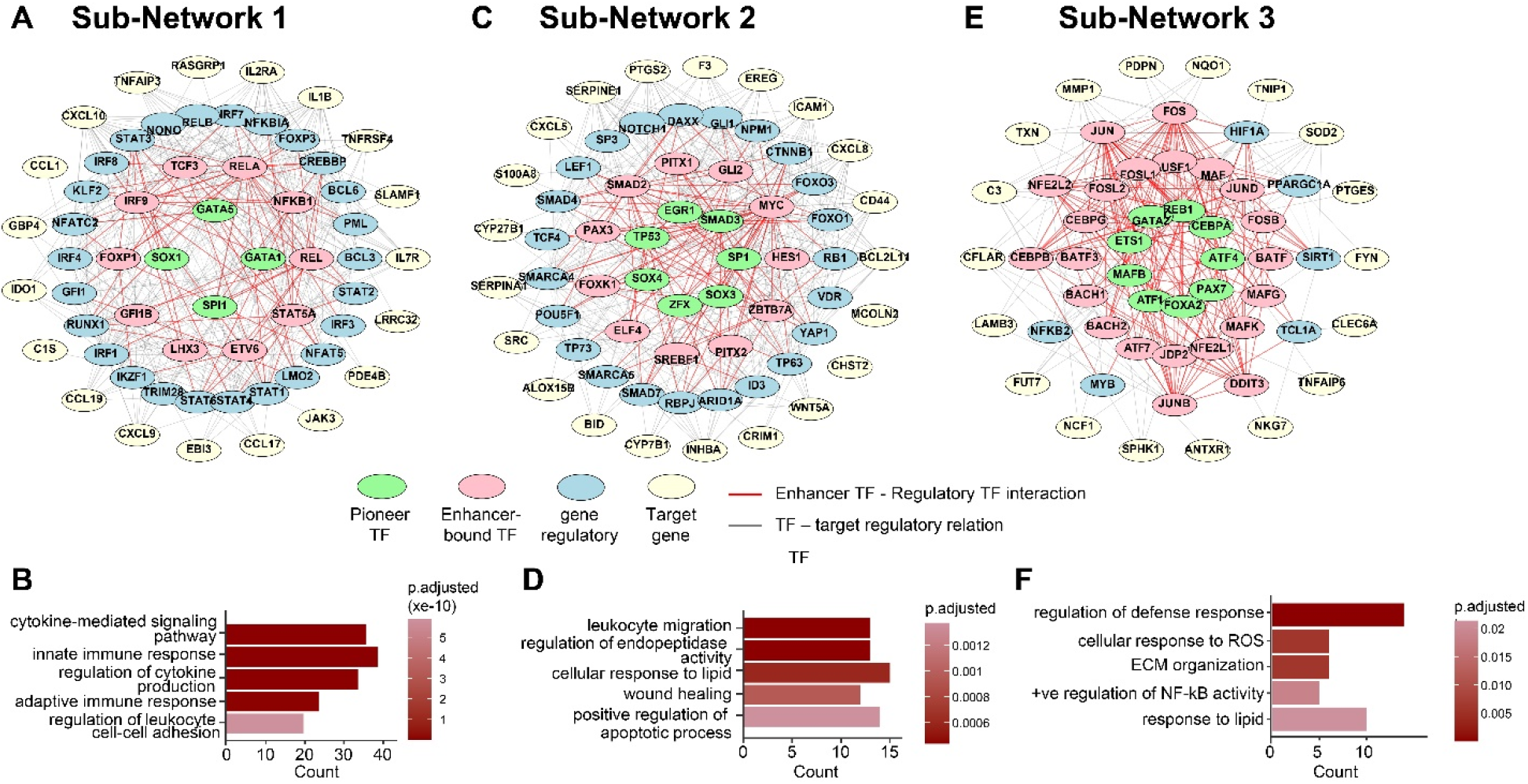
Regulation of macrophage gene expression by enhancer loops that promote interaction between enhancer TFs and TFs at target promoters. **A-F.** Clustering of Pioneer/enhancer TF – regulatory TF – target gene subnetworks (A, C and E) and GO-BP pathways enriched in DEGs in each subnetwork (B, D and F). For each subnetwork, the top TFs and target genes are shown, along with their connectivity. Red lines represent protein-protein interactions obtained from STRING. Grey lines are inferred by regulatory relations from IPA. ROS-reactive oxygen species; ECM-extracellular matrix; +ve-positive.

### 2.8 HT-induced enhancers harbor genetic variants related to metabolic disease

Genetic studies have shown that most disease-associated variants reside in non-coding regions, with many mapping to cis-regulatory elements such as enhancers (*42–44*). To determine whether HT-regulated enhancers in human macrophages contribute to metabolic disease risk, we retrieved genetic variants reported to be associated with metabolic diseases and their complications and examined their presence within HT-regulated enhancer regions (*45*). These included SNPs in key TF genes like *MAF* involved in macrophage polarization (*46*), *SHQ1*, a ribonucleoprotein assembly factor involved in ER stress and ribosome biogenesis (*47*), *TM4SF4*, genetically associated with maturity-onset diabetes of the young (MODY), a monogenic form of diabetes (*48*), *CDKN2B-AS1*, gene for lncRNA ANRIL associated with diabetes complications (*49*), *METAP2P1*, a pseudogene of *METAP2,* which plays a role in fat metabolism and protein synthesis, and is a potential therapeutic target for obesity and T2D (*50*) (**Fig. 8A-F**).

**Fig 8.**
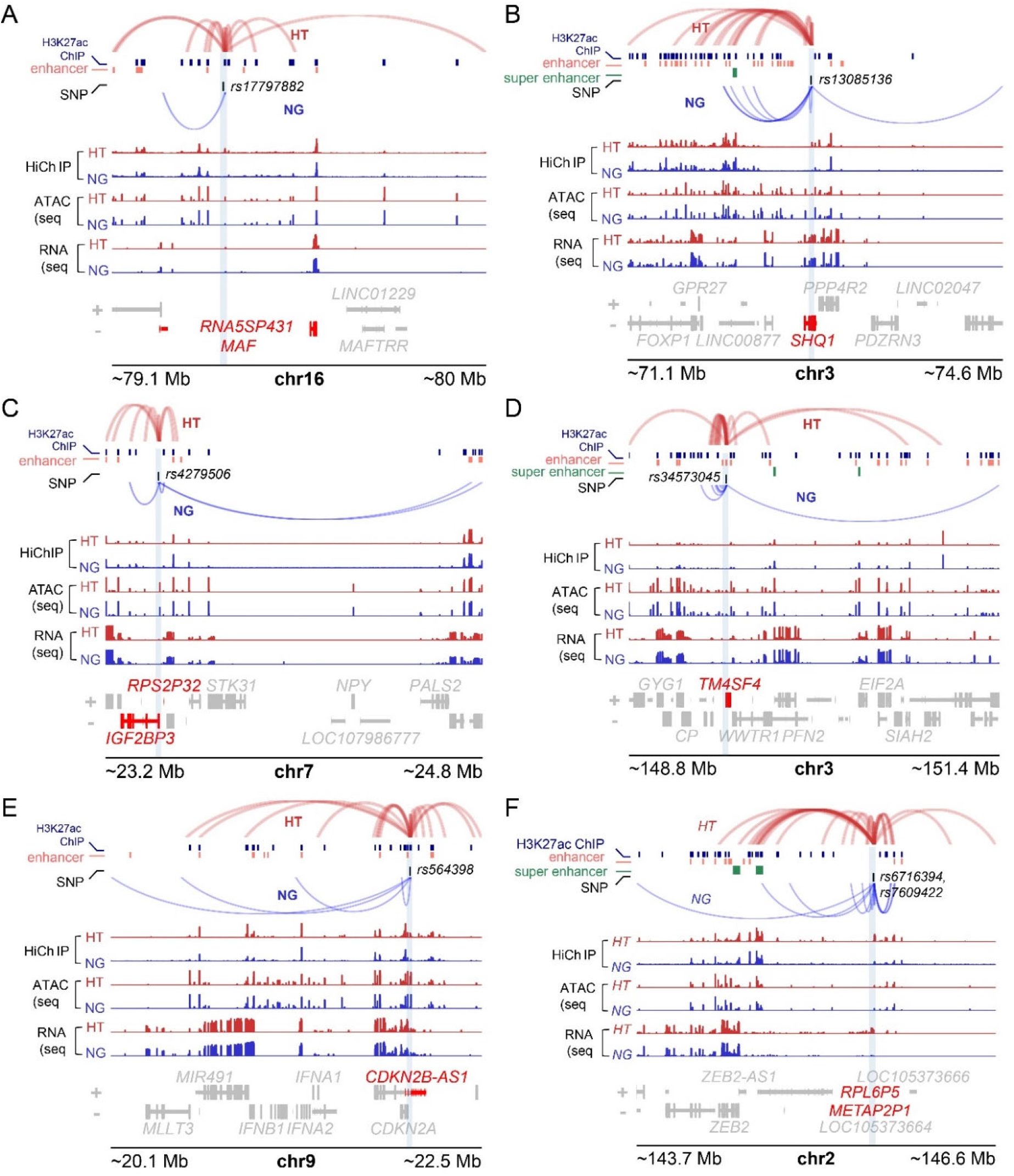
**HT-induced candidate enhancers harbor genetic variations related to T2D**. **A-F**. Genomic tracks from H3K27ac HiChIP and ChIP-seq data showing SNPs associated with differentially regulated chromatin loops between HT (red) vs NG (blue) in human macrophages. Locations of SNPs are indicated by blue color vertical boxes. ATAC-seq (open chromatin), and RNA-seq tracks in human macrophages (this study) and known enhancer and super-enhancer regions in human cells (accessed from the SEdb database) are also shown. The y-axis in all the tracks represents RPKM.

We next assessed whether single nucleotide polymorphisms (SNPs) linked to T1D, T2D and diabetes complications (*45*) reside within HT-regulated enhancer-promoter loops. Indeed, we identified multiple variants within HT regulated enhancer–promoter loops (**Table S3**), almost all of which were reported to be associated with T2D. Specifically, a HT-regulated loop encompassed several variants at the gene locus transcription factor 7-like-2 (*TCF7L2*), which carries the largest genetic risk for T2D (*51*) and is implicated in impaired β cell function, macrophage polarization, and cardiovascular disease (*52*). Furthermore, other loops contained SNPs near genes such as *TNFAIP8, IL7R, PPARG, and FGF6* which are involved in inflammatory response, cholesterol homeostasis, adipogenesis, and insulin sensitivity (*53–56*), core pathways involved in metabolic dysfunction. These results demonstrate that HT-induced enhancers harbor SNPs associated with metabolic dysfunction and could contribute to reshaping 3D chromatin architecture and to altered regulation of genes influencing metabolic diseases, although no overall enrichment of genetic associations were detected compared with all enhancer-promoter loops.

## 3. DISCUSSION

The higher-order chromatin organization shapes the cellular epigenome and enables functional plasticity in response to external stimuli in macrophages (*57*). However, how this controls the dysregulated macrophage phenotypes in diabetes-induced inflammation remains unknown. The current work provides significant advances to the field by demonstrating that HT (mimicking the diabetic milieu) induces coordinated reprogramming of human macrophages across transcriptomic, epigenomic, enhancer connectome and 3D chromatin layers/architecture which is associated with dysregulated macrophage phenotype. We found similar enhancer-promoter interactions in monocytes from diabetic subjects, validated the function of HT induced interactions in regulating key genes associated with inflammation and monocyte/macrophage activation, and identified genetic variants associated with diabetes in these enhancers, thereby providing disease relevance. To our knowledge, this is the first study to comprehensively analyze 3D chromatin remodeling and the enhancer connectome in primary human macrophages under conditions mimicking a pro-inflammatory diabetic milieu. The datasets generated herein can provide a comprehensive and impactful resource for 3D genome remodeling, genome-wide enhancer-loop formation, and TF network hubs associated with macrophage activation in diabetic conditions to advance the field.

Transcriptome profiling showed a shift towards inflammatory signaling and was consistent with a shift away from macrophage alternative activation and homeostatic macrophage functions(*58, 59*). The similarity between HT treated macrophages and primary monocytes from donors with T2D across canonical pathways, functions, and upstream transcription regulators/TFs underscores the disease relevance of this model and supports its utility to dissect diabetic innate immune dysregulation.

ATAC seq data established a clear mechanistic link between the HT transcriptome and chromatin remodeling. Regions gaining accessibility in HT were enriched for inflammatory and innate immune functions and for motifs of TFs that mediate macrophage activation. These data demonstrate that HT augments binding of signal-dependent TFs at pre-existing or newly opened enhancers while promoting condensed (repressed) chromatin at homeostatic loci, thereby priming macrophage state toward sustained inflammation.

H3K27ac HiChIP connectome maps revealed extensive rewiring of enhancer–promoter looping (enhancer connectome) under HT, including numerous differentially regulated interactions. Integrative analysis of Enhancer connectome with ATAC-seq, CHIP-seq, and RNA-seq data revealed that genes with concordant gains in enhancer looping and chromatin accessibility exhibit the largest expression changes, indicating mechanistic coupling between 3D communication and transcriptional output. At typical inflammatory loci (e.g., *IL1B*, *NFKBIA*, *TNIP1*), HT strengthened enhancer loops and increased accessibility, whereas downregulated antioxidant/immune suppressor genes (*SELENOP*, *CD300A*) lost chromatin loops and accessibility. GREAT analysis and motif enrichment at loop anchors implicated inflammatory/immune processes and NF-κB/AP-1 binding, further supporting the central role of HT-regulated enhancers in the inflammatory activation of macrophages.

By overlaying publicly curated enhancer catalogs with our HiChIP enhancer-promoter loops, we identified a novel distal macrophage-specific enhancer ChemDE ∼50 kb upstream of chemokine cluster whose activity and long-range contacts with *CCL2* and other chemokine promoters were selectively enhanced by HT. Furthermore, HT increased H3K27ac at ChemDE, enhanced its interactions with *CCL2*, and induced *CCL2* and *CCL1* expression. CRISPRi targeting of ChemDE significantly blunted HT-induced *CCL2*/*CCL1*, providing a causal link between distal ChemDE enhancer function and chemokine production. Chemokine CCL2 primarily recruits monocytes/macrophages via CCR2, while CCL1 attracts T cells via CCR8. Both are implicated in various inflammatory diabetes complications, such as DKD and CVDs. Similarly, HT enhanced distal enhancer contacts with the *IRAK2* promoter, increased H3K27ac at these elements, and induced *IRAK2* expression. IRAK2 plays a critical role in Toll-like receptor (TLR) signaling by interacting with TLR-associated adaptor proteins, such as MYD88 and MAL, to promote sustained NF-κB activation and cytokine production (*60*). These enhancer–promoter interactions at *IRAK2, IL1B* and *NDRG1* enhancers were also elevated in T2D patient monocytes, supporting human disease relevance. IL1B is a major cytokine that promotes inflammation in diabetes and its complications (*61, 62*), whereas NDRG1 is a multifunctional protein implicated in vascular inflammation, immune response, and hematopoietic stem and progenitor cell differentiation, including the expansion of myeloid-lineage cells (*63, 64*). Together, these data reveal for the first time key enhancer hubs that drive pro-inflammatory cytokines, chemokine and TLR adaptor programs, and critical regulators of vascular inflammation and hematopoietic stem cells under diabetic-like conditions, suggesting signaling pathways by these proteins, as well as their cis-regulatory elements, as potential therapeutic targets.

Furthermore, HT altered A/B compartments and shortened TADs while increasing their number, suggesting disruptions of TAD boundaries at inflammatory loci, such as the *NFKBIA* locus on chr14. These changes provide a structural framework that may facilitate the formation or insulation of specific enhancer–promoter contacts, enhancing stimulus responsiveness at key genes. Although most enhancer–promoter interactions are constrained within TADs, our data indicate for the first time that diabetic condition-like signaling can remodel TAD architecture at relevant loci, likely facilitating rapid and sustained induction of inflammatory gene expression. Thus, HT-induced changes in 3D genome architecture can play an important role in inflammatory gene activation and likely, inflammatory/glycemic memory.

The motif enrichment and network analyses organized the HT response into a hierarchical TF architecture in which pioneer TFs (SPI1/PU.1, C/EBP family, MAF, SOX, FOXO) nucleate enhancer accessibility and 3D microenvironments that recruit signal dependent and inducible TFs (NF κB, AP 1, STATs, IRFs, NRF2/BACH, ETS). Importantly, integration of motif analyses and TF protein-protein interactions revealed three distinct subnetworks controlling cytokine signaling/proliferation, migration/wound healing, and redox/ECM dynamics, which correspond to key inflammatory and monocyte/macrophage activation processes like monocytosis, transmigration, and enhanced cytokine production. The interactions between enhancer-resident pioneer motifs and loop-mediating anchor regions suggest that chromatin looping is integral to the assembly of these TF networks at disease-relevant loci. In addition, diverse phenotypes induced by HT further support the utility of these loops in identifying TF hubs mediating dysregulated macrophage functions in diabetes-related inflammation. Gene regulation by these subnetworks contributes to a low-grade inflammatory state characterized by dysfunctional cytokine signaling, innate immune activation, and dysfunction of leukocytes associated with diabetes and its complications (*62, 65–67*). Thus, our findings provide new mechanistic insights into the key roles of 3D chromatin architecture and TF hub formation driving distinct biological responses in macrophages that can contribute to the inflammation in diabetes.

Notably, we observed HT regulated enhancers harbor SNPs associated with metabolic disease and its complications, including variants at or genes enriched in NF-κB signaling, PI3K signaling, cholesterol homeostasis, and adipogenesis, linking macrophage enhancer activation to pathways central to insulin resistance, dyslipidemia, vascular inflammation, and end organ disease. Our findings can advance mechanistic investigations into how variants in noncoding regions modulate enhancer strength or TF occupancy in contexts in which those enhancers become engaged (e.g., in response to hyperglycemia and inflammatory cytokines), thereby shaping inter-individual risk for chronic inflammation and its complications.

Collectively, our data support a multilayered model of macrophage activation in the diabetic milieu: HT activates signal-dependent TFs and coregulators that bind and acetylate enhancers, leading to chromatin opening and rewiring of long–range enhancer–promoter communication within a remodeled TAD/compartment landscape, culminating in robust induction of inflammatory genes. From a translational standpoint, these insights identify several therapeutic opportunities: (i) chemokine axis blockade (e.g., CCL2/CCR2) (*65*) as shown by ChemDE function, (ii) proximal inflammatory nodes (e.g., inhibitors of IRAK inhibitors and IL-1β ) to dampen TLR/IL-1R and inflammasome pathways (*68*), and (iii) coactivator/acetylation machinery (e.g., BRD4 inhibition) (*69*) to curb H3K27ac-marked enhancer activity and attendant looping. Enhancer-centric pharmacodynamic biomarkers such as H3K27ac or interaction signatures at sentinel loci, may, in the future, enable patient stratification and on-target readouts in trials focused on inflammatory complications of diabetes (e.g., DKD, DR and CVD). Evidence shows that JQ1, a small-molecule BET bromodomain inhibitor that blocks BRD4’s coactivator function, reduces vascular inflammation. JQ1 abolished epigenomic changes at inflammatory super enhancers formed by NF-kB in vascular cells and ameliorated high cholesterol diet-induced atherosclerosis and Angiotensin II induced hypertension, and gene expression in cardiac remodeling/heart failure (*70–72*). It would be worth examining if JQ1 acts by interrupting the 3D chromatin changes under diabetic conditions. In addition, epigenome editing with CRISPR technologies can be leveraged to reprogram enhancers (*73*) to alter macrophage function or determine the role of enhancers harboring SNPs in inflammation and diabetes.

Notably, our study lays the foundation for additional in-depth investigations into genes that are only regulated transcriptionally (i.e., no changes in ATAC-seq), those regulated via pioneer/lineage-dependent TFs, and those by stimulus-dependent TFs, as well as the ratio of genes regulated epigenetically and higher-order chromatin re-organization. While we identified the well-known NF-kB TF as highly activated, we also observed changes in other TFs, such as SPI1, C/EBP, MAF, AP 1, STATs, IRFs, and ETS) based on promoter-enhancer loops, that might have potentially novel functions in macrophage biology. Overall, our integrated multi-omics analysis reveal that HT drives comprehensive reprogramming of the macrophage epigenome and 3D genome, coupling accessible, acetylated enhancers with rewired enhancer–promoter loops and perturbed higher-order topology to enforce key TF like NF-κB/AP1/STAT/IRF-mediated inflammatory gene transcription, macrophage activation and chronic inflammation (**Fig. S13**). Functional validation of novel distal chemokine enhancer (ChemDE) and disease-relevant rewiring at *IRAK2* reveal regulatory elements linking diabetic signaling to pathogenic gene-expression programs. The enrichment of metabolic disease-associated variants within HT engaged enhancers implicates context-dependent enhancer activation in macrophages as a mechanistic nexus connecting metabolic stress to chronic inflammation and end-organ complications in diabetes.

## 4 MATERIALS AND METHODS

### 4.1 Human CD14^+^ monocytes and their differentiation into macrophages

Blood samples were collected from healthy donors and donors with Type 2 diabetes (T2D, HbA1c 6.0 - 6.7%) after obtaining informed consent and in accordance with institutionally approved IRB protocols (IRB 00123 at City of Hope and IRB 300008475 at the University of Alabama, Birmingham). PBMCs were isolated using Ficoll gradient and CD14^+^ monocytes were purified using immunomagnetic CD14 negative selection kits (Stem cells) as described before (*27*). CD14^+^ monocytes were differentiated into macrophages using M-CSF (50 ng/ml,7 days) (*27*). In some experiments, we isolated PBMCs from leukocyte-reduction system cones collected from non-diabetic individuals at Michael Amini Transfusion Medicine Center (City of Hope), then isolated CD14^+^ monocytes and differentiated them into macrophages.

### 4.2 Sample preparation for multi-omics assays

Human macrophages from three healthy donors were treated with high glucose (HG, 25 mM) for 72 h, and in the last 24 h, TNF-α (20 ng/ml) was added to HG samples (HT). Macrophages treated with normal glucose (NG, 5.5 mM) were used as controls. At the end of the treatment, human macrophages treated with HT and NG, were dissociated from plates using Accutase, counted, and divided into multiple fractions for downstream assays. For RNA isolation, cells were immediately lysed in Qiazol and stored frozen at -80 ^0^C. Macrophages used for ChIP-seq and HiChIP were frozen in liquid nitrogen until further use. On the other hand, freshly isolated macrophages were used for ATAC-seq analysis. Results were compared between HT vs NG in all the assays.

### 4.3 RNA-seq Analysis

Total RNA of macrophages from three donors treated separately with NG and HT was isolated using kits from Qiagen or Zymogen and on-column DNase digestion to remove DNA contamination. RNA-seq analysis reads were aligned with human genome Hg38 and differentially expressed genes (DEG)s were identified as described previously (*27, 74*). DEGs were further analyzed to identify enriched GO biological processes, signal transduction pathways and diseases and functions using GSEA and Ingenuity pathway analysis (IPA).

### 4.4 ATAC-seq analysis

ATAC-seq was performed as described (*72*) using macrophages from three separate donors. Sequence reads were aligned to the human genome (Hg38) and peaks in each sample were identified using Genrich. Differentially regulated ATAC-seq peaks representing open chromatin were identified following our bioinformatics pipeline (*75, 76*). ATAC-seq libraries exhibited the anticipated fragment length distribution, with a predominance of short fragments indicative of open chromatin regions between nucleosomes, and a gradual decline in the number of longer fragments corresponding to nucleosome-spanning regions. We identified >71,000 combined peaks with >=100bp commonly found in at least 3 samples. Unique reads within these peak regions were quantile normalized, and differentially regulated peaks were identified (logFC>=1.5, p<0.05, and reads >=50 at least in one sample).

### 4.5 HiChIP assays and data analysis

ChIP-seq and HiChIP assays were performed with NG and HT treated human macrophages described in RNA-seq and ATAC-seq using published protocols (*36, 77*), an H3K27ac antibody and kits from Dovetail Genomics/Cantata Bio. H3K27ac-HiChIP was performed in two replicate samples. In Sample 1, we used macrophage chromatin from a single non-diabetic donor treated with NG and HT. In Sample 2, we used macrophage chromatin pooled from two donors treated with NG and HT, to obtain adequate material needed for HiChIP profiling (due to insufficient material from each donor). H3K27ac-ChIP-seq was performed using pooled chromatin remainder after HiChIP from three non-diabetic donors. HiChIP data for samples 1 and 2 were processed separately using the Dovetail genomics pipeline, as recommended in their online documentation (https://dovetail-analysis.readthedocs.io/en/latest/). In brief, fastq reads were aligned to the human genome Hg38 using BWA-mem (*78*), followed by the identification of valid ligation junctions and removal of PCR duplicates using Pairtools (*79*). The resultant .pairs files were used to generate the contact matrix per sample using Juicer Tools (*80*). Significant loops (5kb resolution, qvalue < 0.05), TADs and A/B compartments were identified using the HiC-DC+ software package (*34*). TADs were identified within the HiC-DC+ framework using the TopDom algorithm (*81*). Differential TAD boundaries between HT and NG were detected using the software TADCompare (*82*). Chromatin regulatory loops identified in our study were compared with public Gen-Hancer datasets from multiple cell types (*83*) to identify novel enhancer-promoter interactions in human macrophages under diabetic conditions. . Integration of RNA-seq/ATAC-seq/ChIP-seq/HiChIP was performed using R v4.5 and Bioconductor (*84*). Enriched GO Biological processes and signaling pathways were identified using GSEA and Ingenuity pathway analysis (IPA). Transcription factor networks were identified after integration of Upstream regulatory analysis in IPA, STRING database of protein-protein interactions and gene expression analysis. Genomic plots involving RNA-seq and ATAC-seq were created using the ggplot2 package (*85*) within the R framework, version 4.3. Genomic tracks and contact heatmaps showing HiCHIP loops were created using the R package PlotGardener (*86*).

### 4.6 RT-qPCR and ChIP-qPCR

Gene expression was analyzed using RT-qPCR of total RNA as described before, using gene-specific primers (*27, 87*) . Gene expressions were analyzed using 2^-ΔΔT^ after normalization with internal controls *RPLP0* or *HPRT.* ChIP assays with H3K27ac antibodies were performed as described earlier (*27, 87*) and ChIP DNA was analyzed by QPCR using the indicated primers. Data was normalized with input and results expressed as %Input. Primers used are listed in Supplementary **Table S4**.

### 4.7 Chromatin Conformation Capture (3C) assays

These assays were performed using published protocols (*84*). Primary human macrophages treated with NG or HT, or those derived from volunteers with documented T2D and healthy controls (3-5 million cells) were fixed with 4% formaldehyde, lysed in HiC buffer, and chromatin was digested with HindIII Hi-fidelity, followed by dilution and ligation overnight as described (*18*). Then, crosslinks were reversed, and DNA was purified using phenol/chloroform extraction and ethanol precipitation. The purified DNA was analyzed by quantitative PCR (qPCR) using primers targeting HindIII sites in chromatin loops, as determined by HiChIP results. Primers used are listed in Supplementary **Table S4.**

### 4.8 CRISPR interference (CRISPRi) at enhancers

CRISPRi of HT-induced enhancers was performed by co-transfecting dCas9-SALL1-SDS3 mRNA (*39*) with custom-designed guide RNAs (sgRNA)s targeting HT-induced enhancers or commercially available control non-targeting sgRNA and *PPIB* sgRNA (Horizon). The custom enhancer sgRNAs were designed using the publicly available CHOPCHOP web interface (*88*), utilizing chromosome coordinates in default settings. THP1 monocytes (5 million) were electroporated with a mixture of CRISPRmod CRISPRi EGFP dCas9-SALL1-SDS3 mRNA (5 ug, Horizon Discovery, Cat# CAS12225) and indicated sgRNAs (5 uM, IDT, Table S4) and RNase inhibitor (1µl of 40U/uL, Roche Cat# 03335402001) using the Maxcyte platform. The transfected cells were allowed to recover in complete medium for 4 h, then plated in medium containing PMA (20 ng/mL) for macrophage differentiation. After 24 h, resultant THP1 macrophages were treated with NG or HT, and total RNA was extracted for RT-qPCR analysis of gene expression. In some experiments, we also performed CRISPRi in macrophages derived from CD14^+^ monocytes from nondiabetic donors, using methods as described above for THP1 macrophages.

### 4.9 Statistical analysis

Results of RT-qPCR, ChIP-qPCR and 3C assays are shown as the mean±SD of ≥3 samples, expressed as fold change over controls. We performed statistical analyses in GraphPad Prism using an unpaired two-tailed t-test (two-group comparisons), Multiple t-tests, or one-way ANOVA with multiple comparisons, as specified in the figure legends. Statistical significance was set at *p*<0.05.

### 4.10 Analysis of TF networks

Enriched TF motifs at differentially regulated gene promoters (Promoter motif, Fig. S13B) were derived via HOMER(*89*), using the human genome as background. To identify TF motifs enriched in enhancer loops (Enhancer-loop motif, Fig. S13B), we gathered all differentially expressed HiCHIP loop anchors (|logFC| > ±1) overlapping with annotated enhancer regions in the human genome (using R package Annotatr (*90*)) and performed enrichment analysis in HOMER. In each case, an adjusted pvalue of 0.1 was used as the significance threshold. In addition, predicted TFs with differential activity in HT vs NG, as predicted by IPA based on gene expression changes (Predicted TF, IPA), were obtained using the following criteria: z-score > ±0.5; Overlap pvalue < 10^-5^; Target count > 15. For constructing the TF-target network, we focused on the superset of TFs constituting the Enhancer-loop motifs and IPA predicted TFs. To minimize false positives, the TFs that were identified among the promoter-motifs, but absent among the IPA predicted and the Enhancer-loop TFs were excluded. Protein interaction data (physical links) was downloaded from the STRING repository (*91*). To construct the HT specific interaction network, edges where both nodes originate from enhancer motif TFs, IPA predicted TFs or the differentially expressed genes (|logFC| > ±1; padj < 0.1) were retained. The resulting TF-target network (consisting of 129 TFs and 530 target genes) was clustered into three distinct communities (with 124, 105 and 53 differentially expressed target genes respectively and zero overlaps) based on the leading eigenvector of the graph modularity matrix, as implemented in the cluster_leading_eigen() function of the igraph R package (*92*). Enriched GO-BP pathways in each community were identified using over-enrichment analysis, as implemented by the enrichGO() function from the R package ClusterProfiler (*93*).

## AUTHOR CONTRIBUTIONS

Conceptualization: M.A.R., S.B., and R.N. Methodology: M.A.R., S.B., V.M, V.T. and X.W. Validation: M.A.R., S.B., V.M., V.T., and L.Z. Formal analysis: M.A.R., S.B., V.T., V.M., Z.C., and X.W. Investigation: M.A.R., V.M., V.T., L.Z., and L. L . Resources: M.B.G., Data curation: M.A.R., S.B., V.M., and V.T. Writing-original draft: M.A.R., S.B., and R.N. Writing—review and editing: V.M., V.T., Z.C., J.T., Z.B.C, and M.B.G. Visualization: M.A.R. and S.B.; Supervision: M.A.R., R.N., and Z.B.C. Project administration: M.A.R. and R.N. Funding acquisition: M.A.R., M.B.G., Z.B.C. and R.N.

## ACKNOWLEDGMENTS

We are grateful to Dr. Jinhui Wang (Integrative Genomics Core, City of Hope) for assistance with the sequencing. Research reported in this publication includes work performed in the Integrative Genomics Core and DNA/RNA Synthesis Core (supported by the National Cancer Institute of the NIH under grant no. P30CA033572) at City of Hope and Michael Amini Transfusion Medicine Center (City of Hope) for providing leukocyte-reduction system cones.

## FUNDING

This study was supported by grants from the National Institutes of Health (NIH) R01DK143577 (to RN and ZBC); R01 DK065073 and R01 DK081705 (to RN), R01 HL106089 (to RN and ZBC), NIH grant R35HL171550 (to ZBC), R01EY012601, R01 033620, EY038525 (to MBG), a City of Hope Excellence award (to RN), the George & Irina Schaeffer Foundation (to RN), and an AR-DMRI innovative grant at the City of Hope (to MAR and RN).

## Conflicts of Interest

The authors declare no conflict of interest.

## Data Availability Statement

Raw RNA-seq, ATAC-seq, ChIP-seq, and HiCHIP data generated in this study have been deposited in the GEO database and are available under accession numbers GSE334742, GSE334739, GSE334740, and GSE334746, respectively.

## Notes

### Competing Interest Statement

The authors have declared no competing interest.

### Summary of Updates

The revised manuscript strengthens human disease relevance and reproducibility. Revisions clarify interpretation, methods, validation, sample descriptions and sizes, and analyses. The A/B compartment analysis was removed because HiChIP contact matrices may underrepresent B compartments, while public CTCF ChIP-seq data and an additional HiChIP sample supported TAD boundaries and enhancer-connectome alterations. The revision also clarifies Chemokine distal enhancer (ChemDE) visualization, provides validation of ChemDE activity in primary human CD14+ monocyte-derived macrophages, expands transcription-factor analysis criteria, improves figure resolution, and reduces the figure count from 10 to 8 by moving two figures to supplementary materials.

