## Supplementary material for "Epigenome Alterations and 3D Chromatin Architecture Remodeling in Inflammatory Macrophage Activation under Diabetic Conditions": Supplemntal Material

**This PDF includes:**

Figs. S1-S13

Tables S1 to S4

**Figure S1: Inflammatory gene regulation and IPA analysis of HT-regulated genes in human macrophages.**

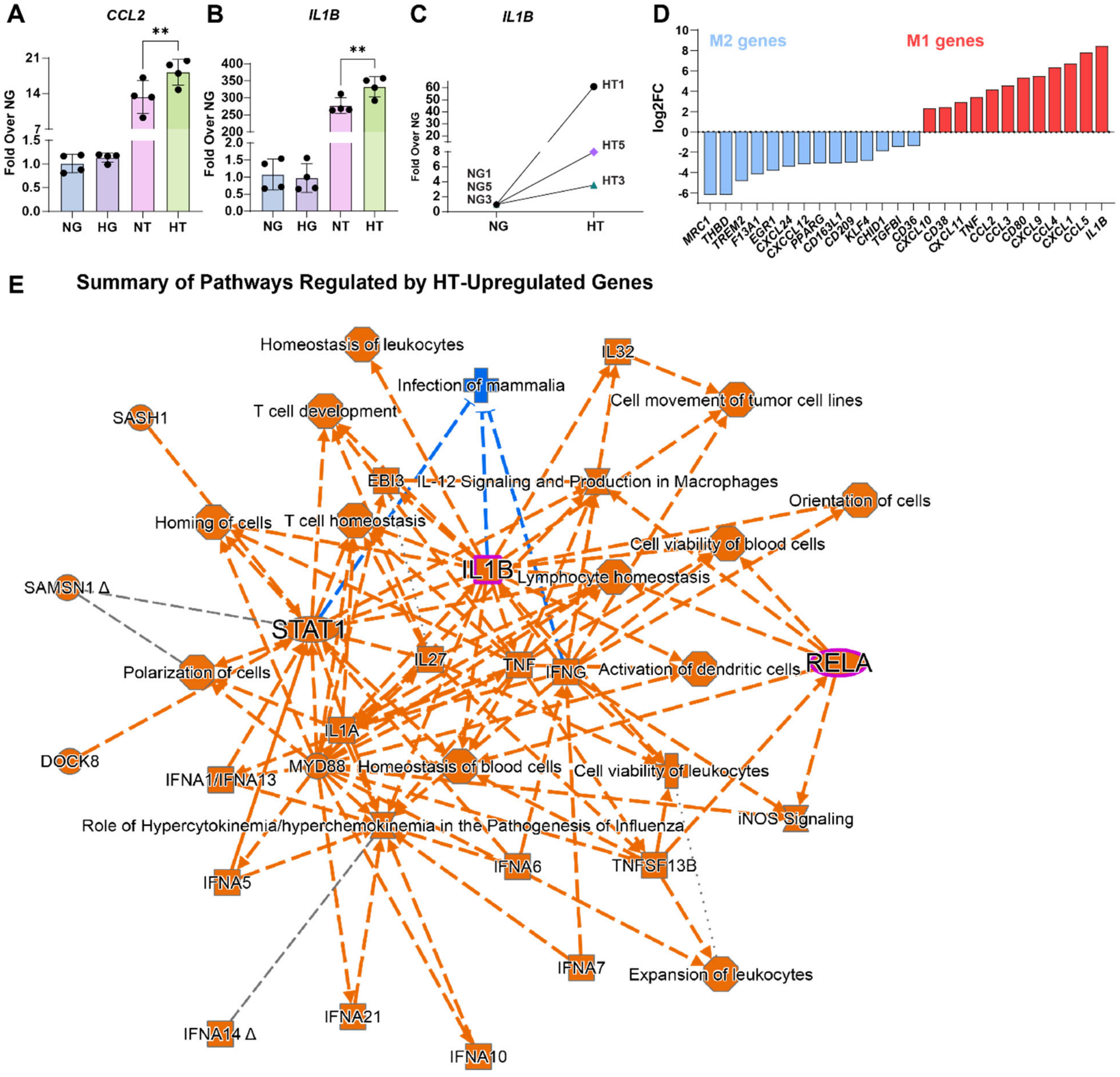


**A-B.** Enhanced inflammatory gene expression by HG + TNF (HT) versus normal glucose +TNF (NT). RT-qPCR analysis of *IL1B* and *CCL2* expression in human CD14$^+$ macrophages from a healthy donor treated with NG, HG, NT or HT. Results were expressed as Fold over NG. **, p<0.01 as determined by Ordinary one-way ANOVA and Sidak's multiple test (n=3 replicates from one donor). **C.** Before-and-after plots showing results of RT-qPCR analysis of *IL1B* expression in human CD14$^+$ macrophage samples treated with HG+TNF (HT) vs NG. CD14+ monocytes from 3 healthy human volunteers were differentiated into macrophages and treated separately with normal glucose (NG) or high glucose (HG) for 72 h; in the last 24 h, TNF-α (20 ng/ml) was added to the HG samples (HT) and gene expression analyzed. Results are expressed as fold over respective control NG treated cells. These samples (NG1/HT1, NG3/HT3, and NG5/HT5) were used in Multi-Omics analyses, including RNA-seq, ATAC-seq (all n=3), H3K27ac-HiChIP (n=2, Sample 1 -single donor and Sample 2-pooled chromatin from two donors), and H3K27ac-ChIP-seq (pooled chromatin from all three samples). **D.** HT treated macrophages exhibited an increased inflammatory phenotype. RNA samples from panel C were analyzed by RNA-seq analysis. Bar graphs show differentially expressed M1 and M2 phenotype genes (log2FC) in HT vs NG (FDR<0.05, n=3). **E.** Summary of signaling pathways enriched in HT-Upregulated genes as determined by Ingenuity Pathway Analysis (IPA).

**Figure S2: Upstream Regulator Analysis of genes upregulated by HT in human macrophages.**

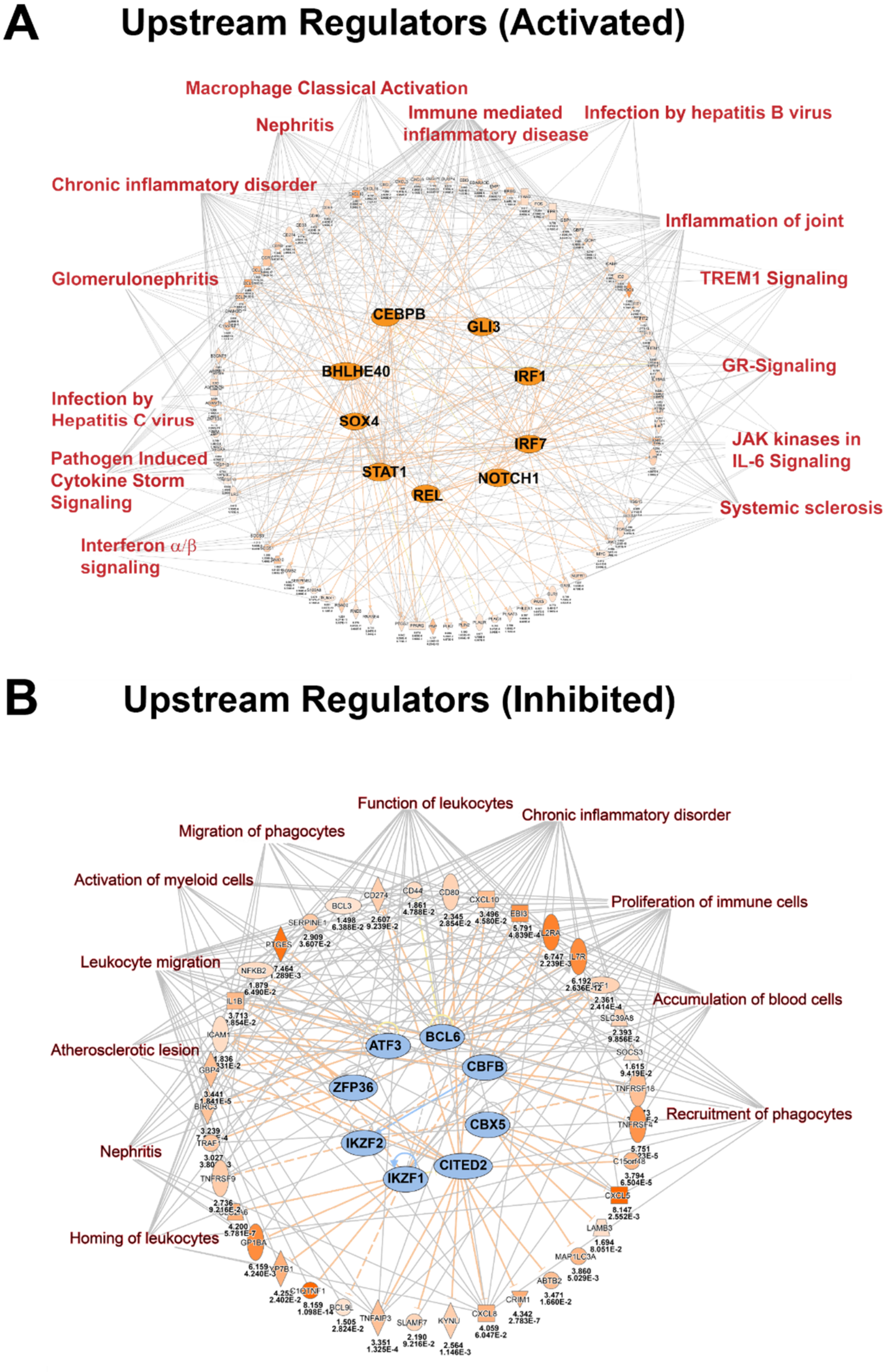


**A-B.** Results of Upstream Regulator Analysis showing transcription factors (TF) activated (**A**) and inhibited (**B**) by HT, and their roles in diseases and functions. Upstream Regulator Analysis of upregulated genes (HT vs NG) in IPA was performed using the default settings.

**Figure S3: IPA analysis of genes downregulated by HT in human macrophages.**

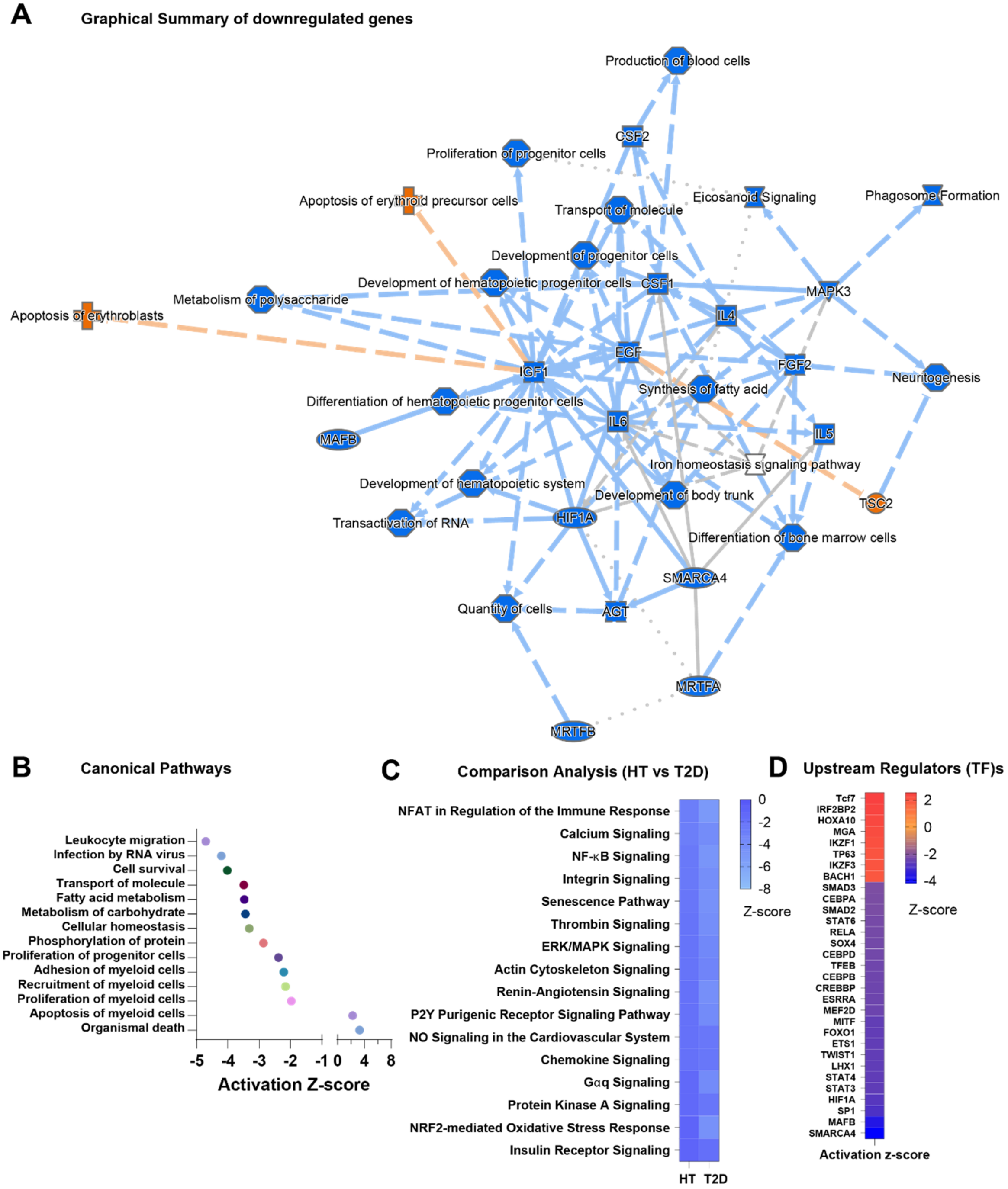


**A.** Summary of pathways and functions enriched in genes downregulated by HT in human macrophages. **B.** Canonical Pathways, **C.** Shared pathways between HT and human T2D (Comparison analysis), and **D**. Upstream regulator TFs, in genes downregulated by HT. IPA of downregulated genes (HT vs NG), including Upstream Regulator Analysis and Comparison Analysis with T2D (vs Controls), were performed using default settings.

**Fig. S4. Results of GREAT analysis and TF Motif Analysis of Downregulated ATAC-seq peaks in human macrophages treated with HT and NG.**

A

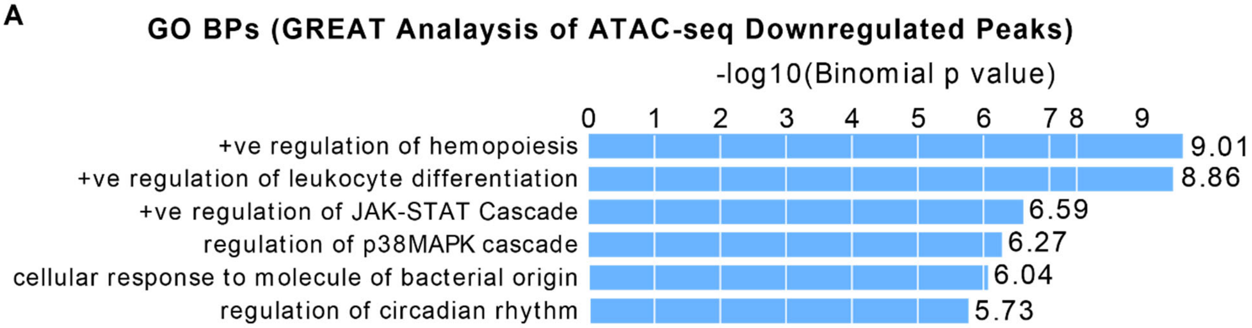


B TF Motifs enriched in ATAC-seq downregulated peaks

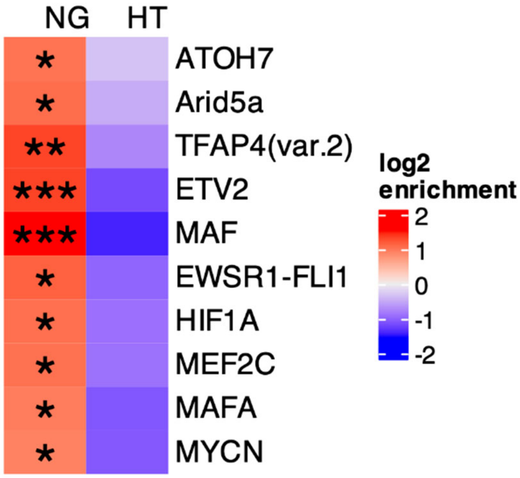


**A.** Results of GREAT analysis showing GO biological processes (BP)s enriched in downregulated open chromatin (ATAC-seq peaks) in HT versus NG treated human macrophages. **B.** Transcription factor (TF) motifs enriched in differentially regulated (downregulated in HT) open chromatin. Only TF-motifs that are shared between RNA-seq and HiChIP are shown. p.adj: *p<0.1-0.01; **p<0.01-0.001; ***p<0.001-0.0001.

**Fig S5. Confirmation of HT-indued changes in TADs and Enhancer connectome in human macrophages in replicate sample.**

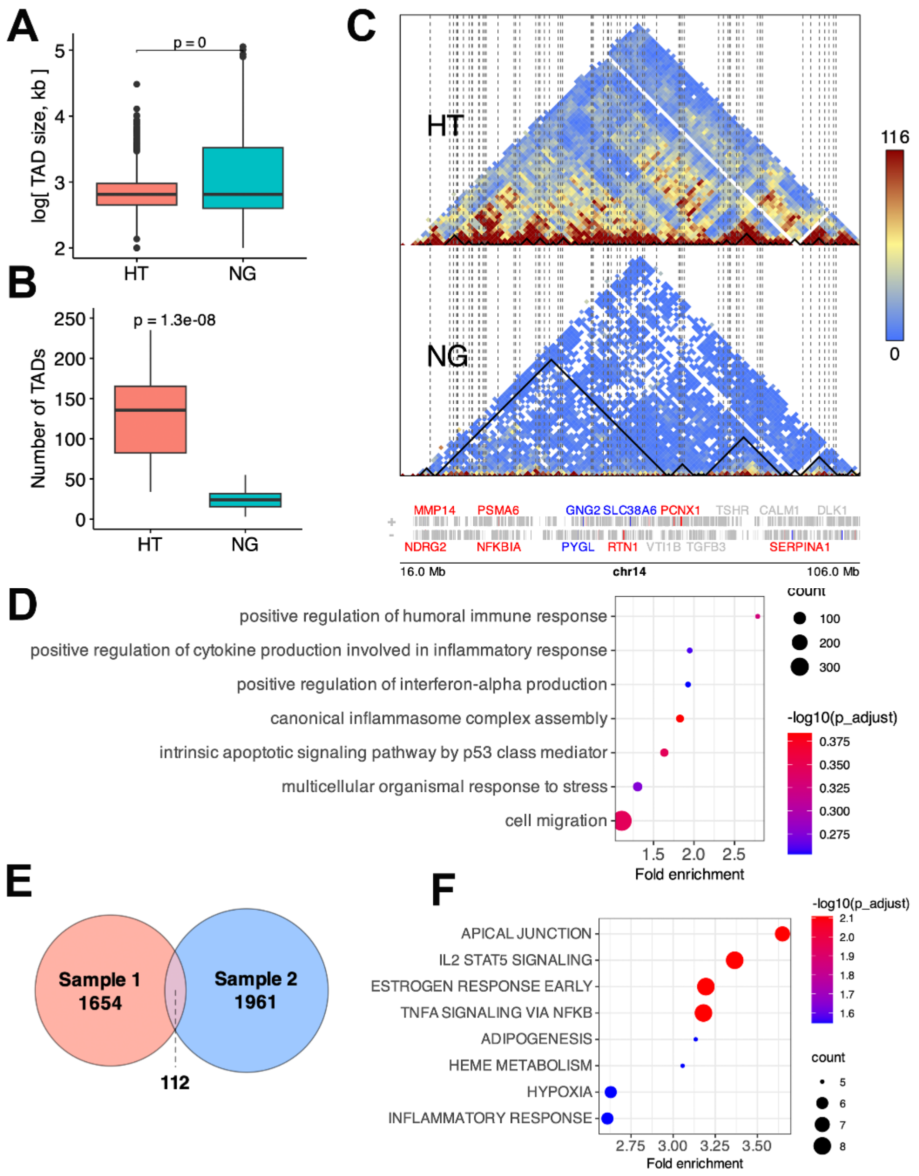


**A-D. Results of TAD analysis performed on replicate macrophage Sample 2, in which H3K27ac HiChIP was performed using pooled chromatin from two donors.** Comparison of TAD sizes (A) and number of TADs (B) between HT and NG in Sample 2. **C.** Heatmap of normalized HiChIP read density in HT and NG on Chr14 for Sample 2. TADs are highlighted by black triangles and disrupted TAD boundaries by dashed vertical lines. HT-regulated genes are highlighted in red (up) and blue (down) colors**. D.** Enriched GO-BPs at disrupted TAD boundaries in HT in Sample 2. **E.** Overlap of HT-disrupted TAD boundaries between Samples 1 and 2. **F.** Enriched pathways (Hallmark) at the common disrupted TAD boundaries between Samples 1 and 2.

**Fig S6. Confirmation of TAD boundaries using CTCF binding sites.**

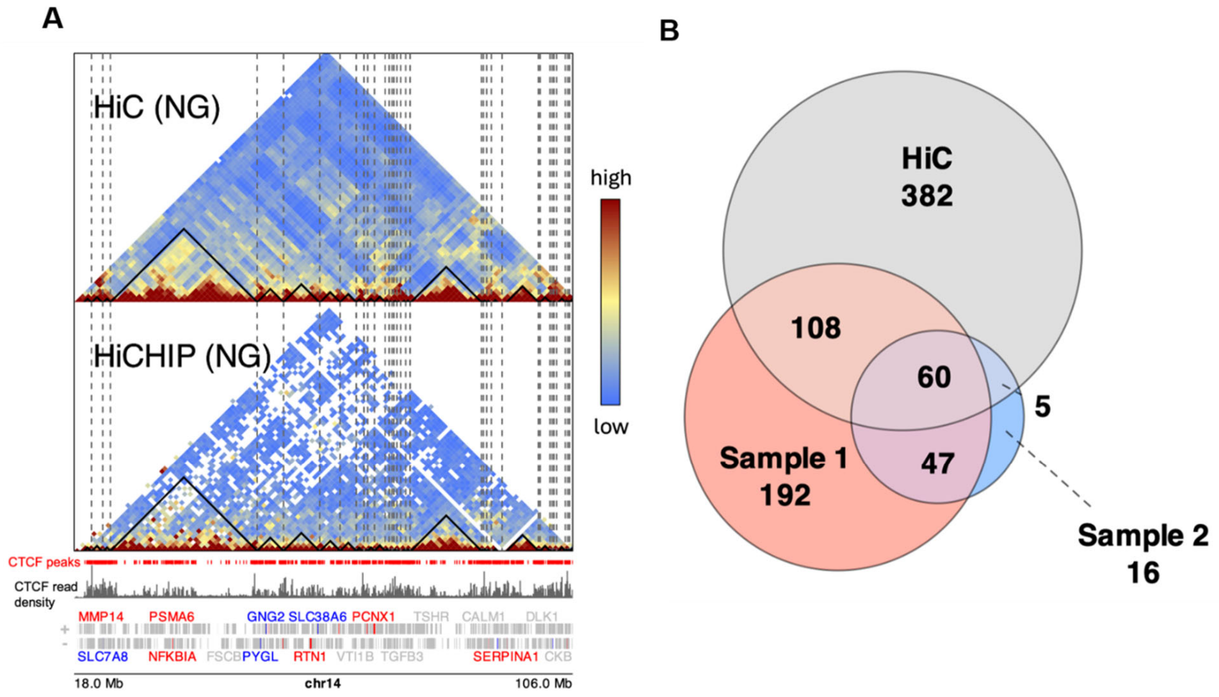


**A.** Heatmap of normalized read density of Chr14 from a publicly available HiC dataset of human M0 macrophages (Array Express, MTAB-10848) compared with that from HiChIP of NG-treated macrophages in Sample 1 in this study. TADs derived from sample 1-NG are highlighted by black triangles and TAD boundaries by dashed vertical lines. CTCF peaks and read density from the same sample used in HiC are shown in red and grey respectively. HT-regulated genes are highlighted in red (up) and blue (down) colors. **B**. Overlap of TAD boundaries among HiChIP of Sample 1-NG, replicate Sample 2-NG and HiC-M0 macrophage.

**Fig. S7. Results of HiChIP analysis in human macrophages treated with HT and NG.**

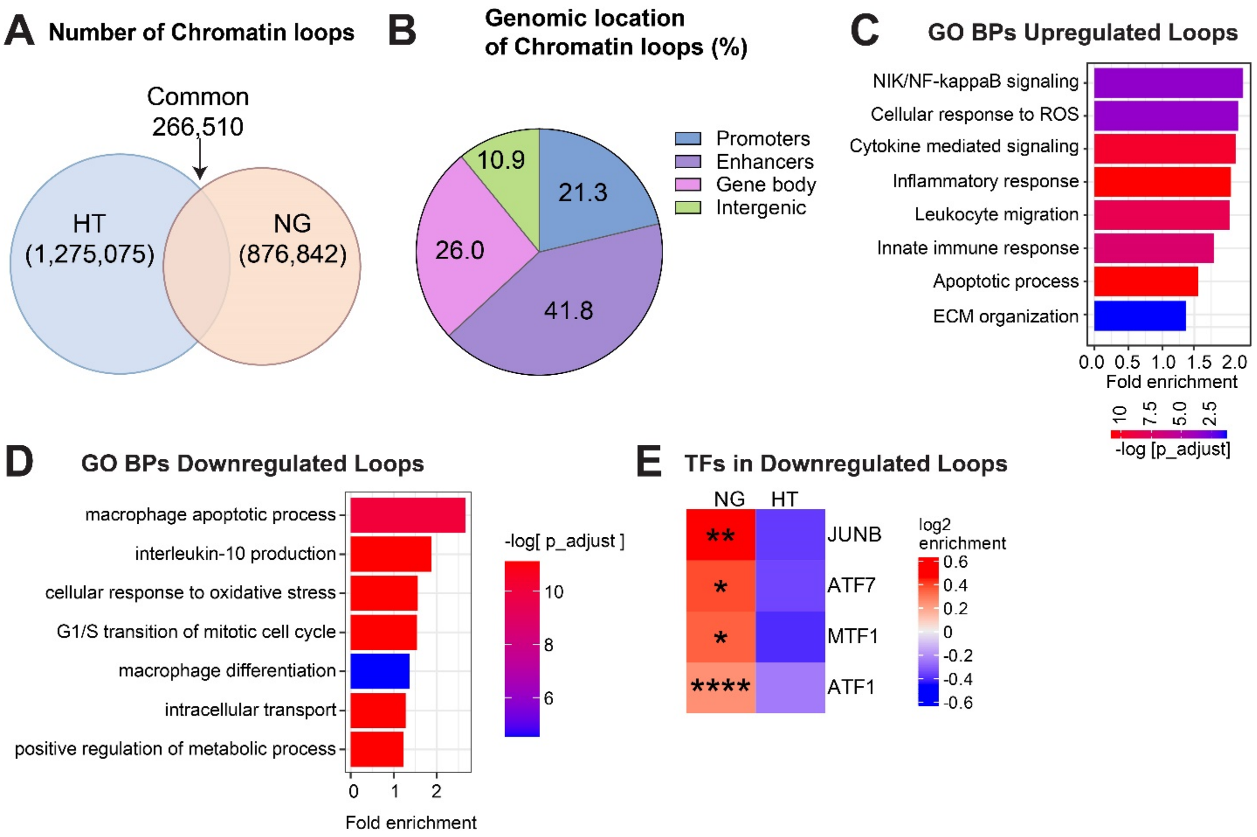


**A-B.** Venn diagram shows chromatin loops in NG and HT samples (**A**) and pie chart shows their genomic distribution (**B**) in human macrophages. HiChIP with H3K27ac antibody was performed in human $CD14^+$ monocytes (from a single donor) differentiated into macrophages and treated with NG or HT. **C-D.** GO BPs enriched in HiChIP loops upregulated (**C**) and downregulated (**D**) in HT vs NG (GREAT analysis). **E.** TF motifs enriched in HiChIP loops (downregulated) in HT versus NG treated macrophages. p.adj: *p<0.1-0.01; **p<0.01-0.001; ****p< 0.0001(hypergeometric test).

**Fig. S8. Visualization of chromatin loops (at indicated loci) mediating enhancer-promoter interactions regulated by HT versus NG in Human macrophages.**

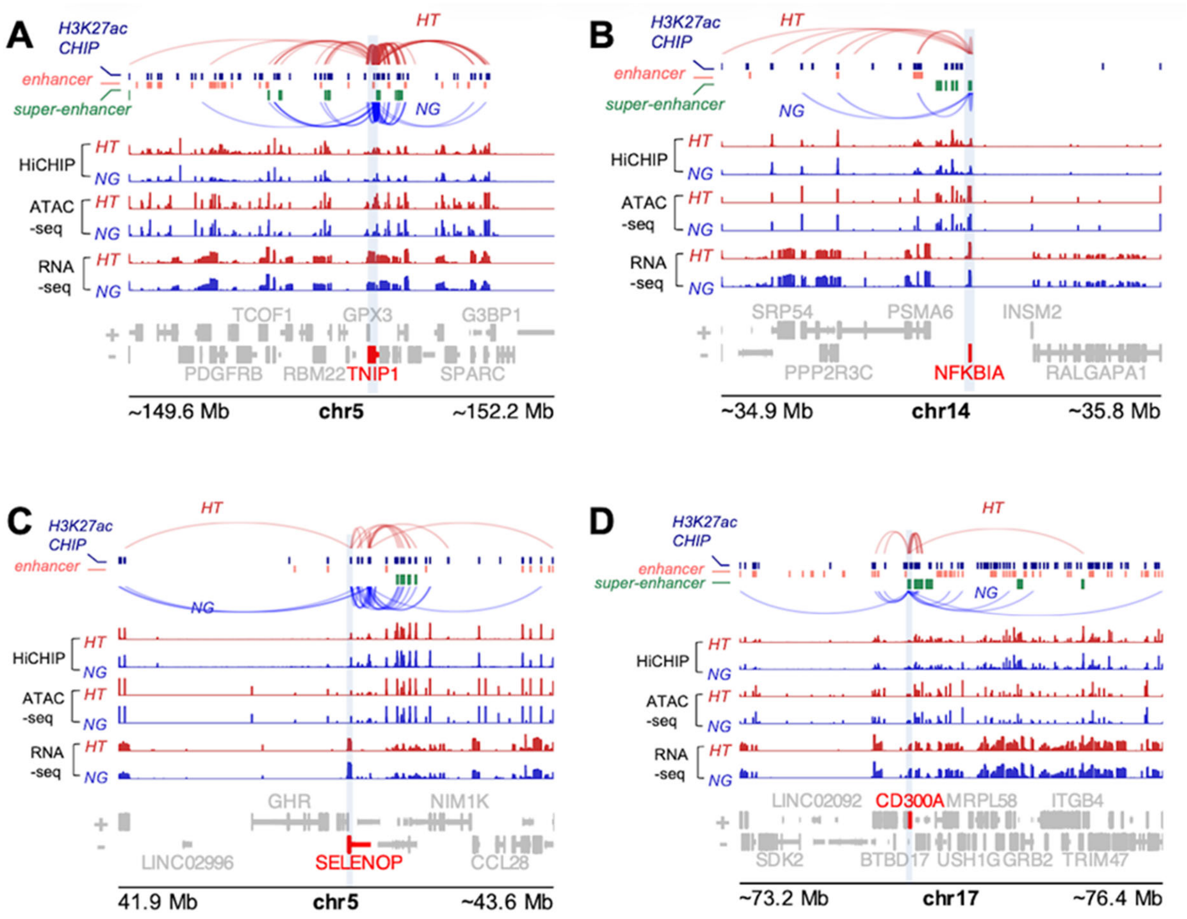


**A-D**. Genomic views of HiChIP tracks showing enhancer-promoter chromatin loops at indicated inflammatory genes (red font text) in HT (maroon color loops) versus NG (blue color loops) -treated human macrophages. The genomic tracks of open chromatin (ATAC-seq peaks) and gene expression (RNA-seq data) have also been shown.

**Figure S9**. **Interaction of chromatin loops between enhancers and promoters of genes associated with inflammatory signaling pathways on the indicated chromosomes.**

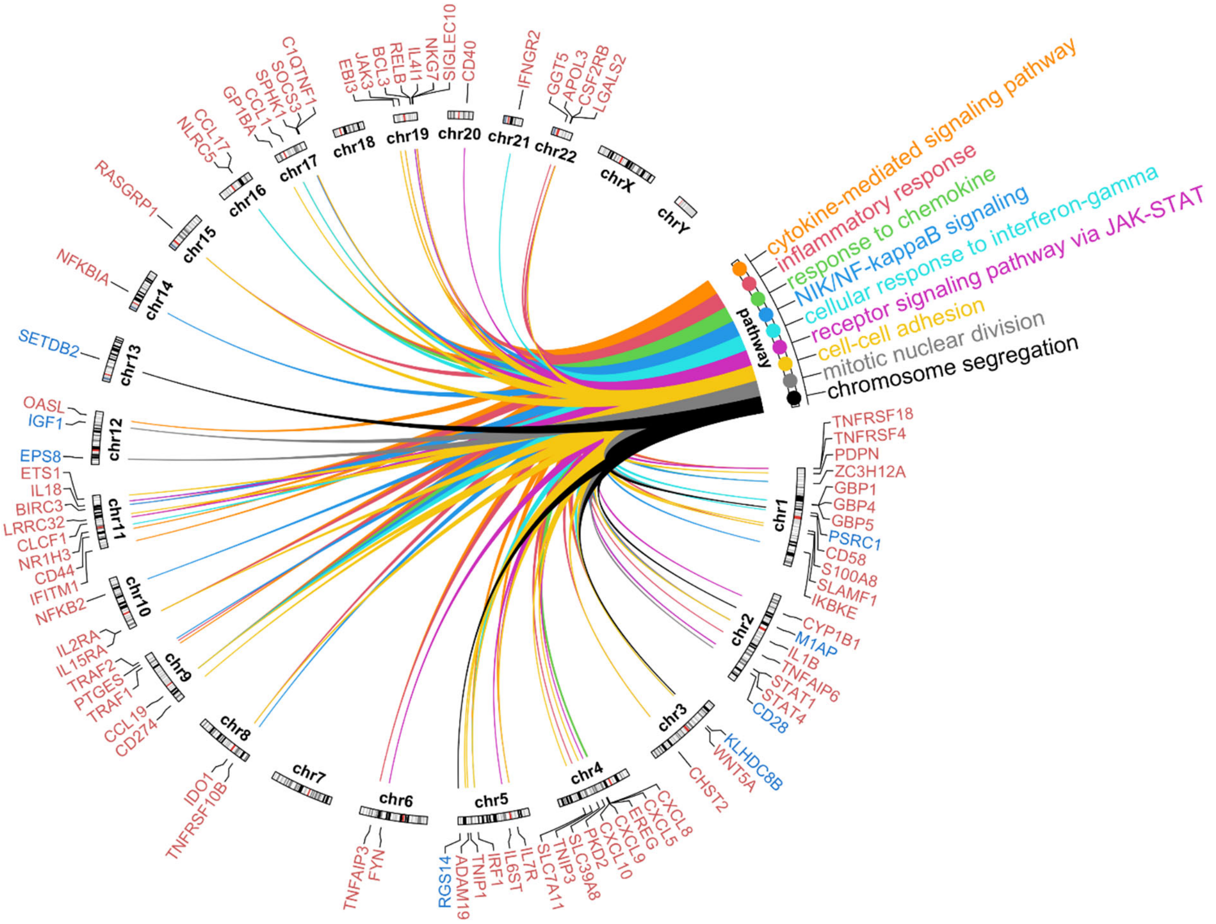


HT-induced chromatin loops (vs NG) in CD14$^{+}$ macrophages were identified using HiChIP analysis with H3K27ac antibody. HiChIP data was integrated with differentially expressed genes (HT vs NG) identified using RNA-seq in macrophages. A CIRCOS plot was generated showing genes from the indicated signaling pathways on different chromosomes (chr).

**Fig S10. Validation of HT-indued changes in Enhancer connectome in replicate sample 2.**

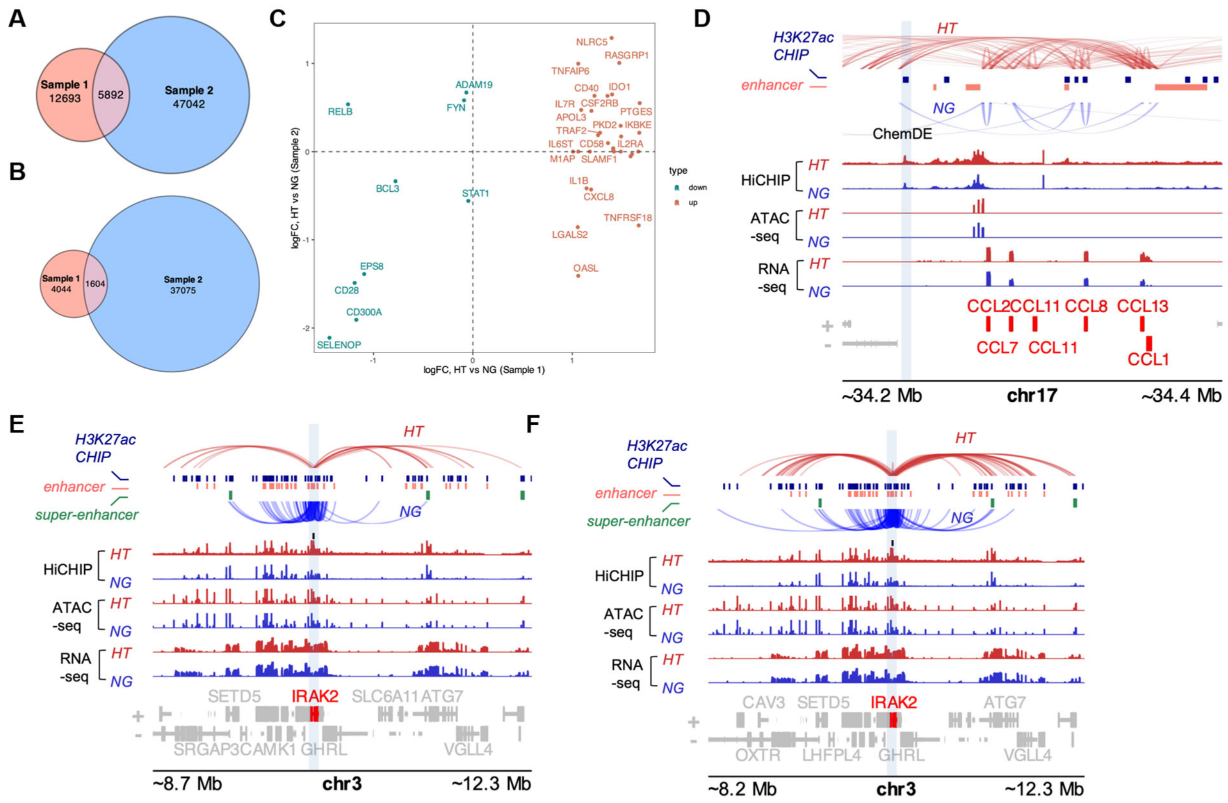


**A-B.** Overlap of differential HiCHIP loops between Sample 1 and replicate Sample 2. Venn diagrams show overlapping HiChIP loops Up (A) and Down (B) in HT vs NG between Sample 1 and replicate Sample 2. **C**. Average log fold change in promoter loops at important gene loci compared between Samples 1 and 2. Genes showing increased and decreased promoter loops in HT in Sample 1 are colored red and blue respectively. **D-F.** Genomic maps showing increased interaction of chromatin loops in HT treated replicate Sample 2 vs NG. **D.** Chemokine distal enhancers (including ChemDE) and promoters of chemokine genes on Chromosome 17. **E-F.** Distal enhancers interacting with the *IRAK2* promoter. H3K27ac ChIP-seq, ATAC-seq and RNA-seq data tracks have also been shown. **E shows** only loops interacting with *IRAK2* promoter bound. **F** shows all loops originating from the *IRAK2* locus, including promoter and gene body regions.

**Fig. S11. 3C assays in macrophages from T2D subjects versus controls.**

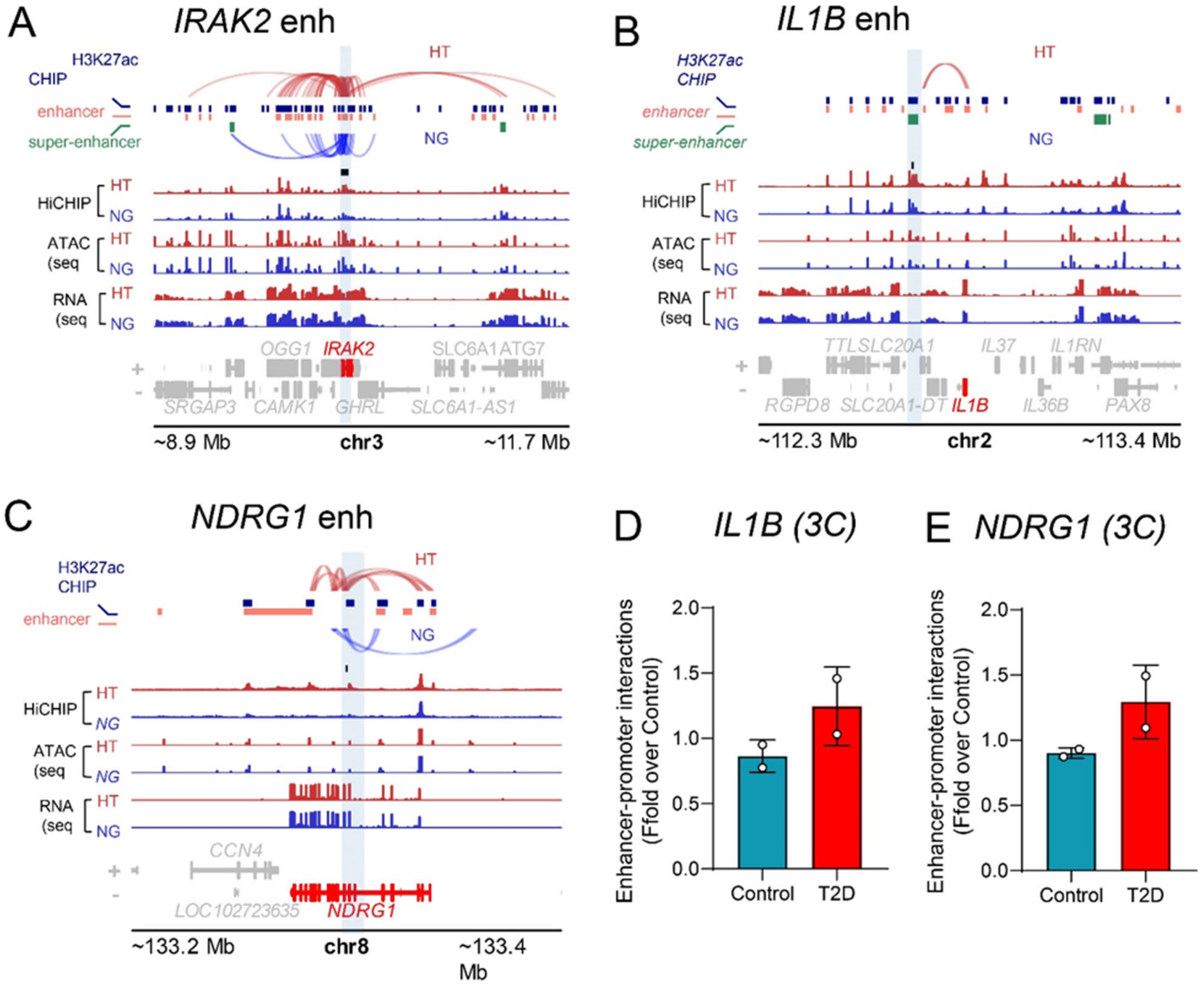


**A-C.** Visualization of HT induced loops mediating enhancer-promoter interactions at the indicated candidate gene loci that were validated using 3C assays in control versus T2D donor cells. The vertical blue color bar indicates the enhancer used in 3C assays. **D-E.** Enhancer-promoter interactions in macrophages differentiated using primary CD14$^{+}$ monocytes from T2D patients versus controls as determined by 3C assays (n=2/group). *IRAK2* loci genomic interactions 3C and related assays are shown in the main manuscript (**Fig. 5G-J**). Candidate gene loci shown were chosen because they showed similar loop alterations in HT treated cells.

**Fig S12. Integrated analysis of TFs enriched in enhancers, open chromatin and promoters under diabetic conditions in human macrophages.**

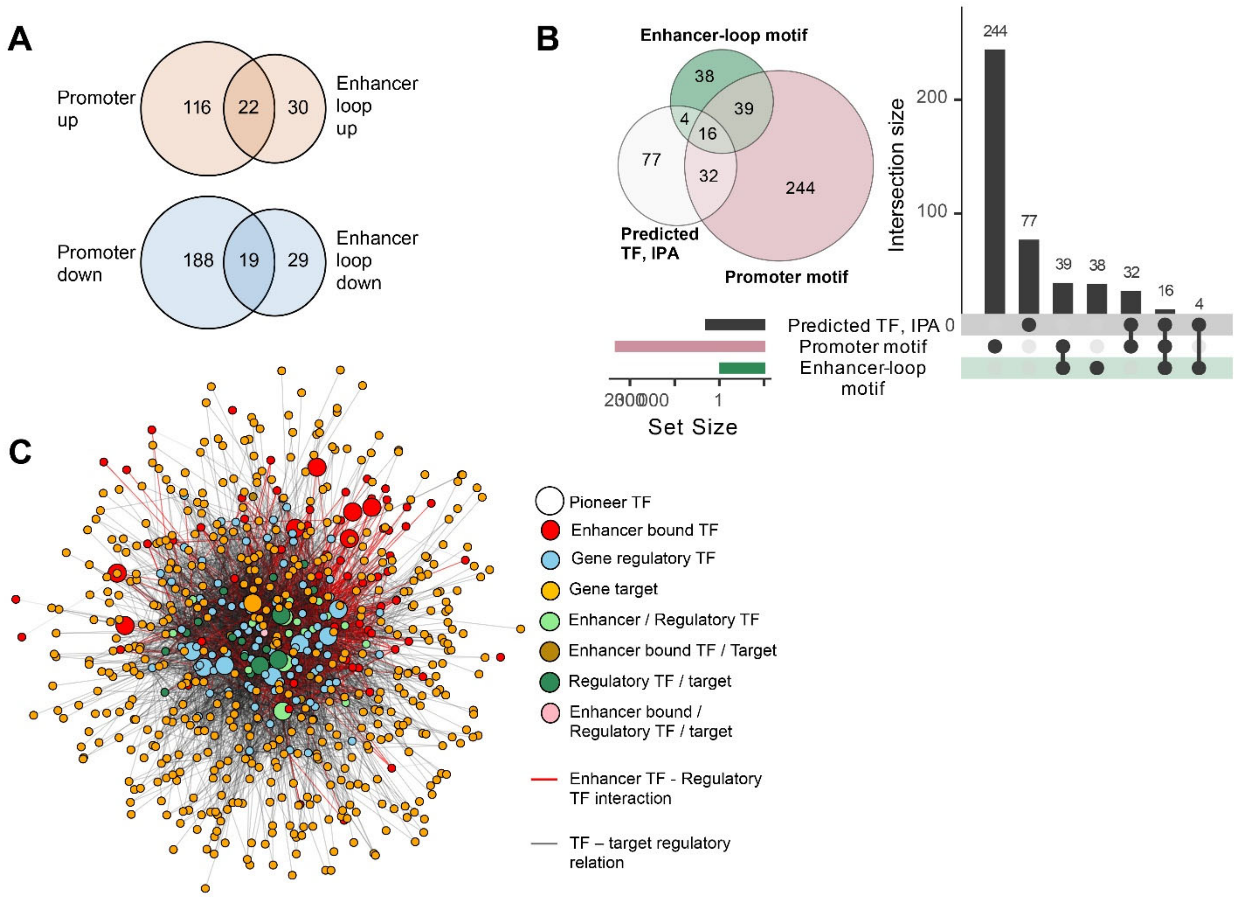


**A.** Overlap between TFs enriched among DNA motifs from promoter regions of DEGs and those from enhancer-associated loops. Up and downregulated sets are separately compared (for the list of TFs, please see **Table S1-S2**.). **B.** Venn diagram and UPSet plot showing overlap among TFs identified through three different analyses, promoter motifs, enhancer loop motifs and IPA. **C.** Inferred network of Pioneer/enhancer TF – regulatory TF – target gene network of upregulated DEGs by combining IPA with known protein-protein interactions from STRING. TFs belonging to various categories are colored as described in the legend. Larger circles represent pioneer TFs. Physical interactions between enhancer-bound and regulatory TFs are highlighted in red.

**Fig. S13. Epigenetic mechanisms and 3D chromatin architecture remodeling in macrophage activation under diabetic conditions.**

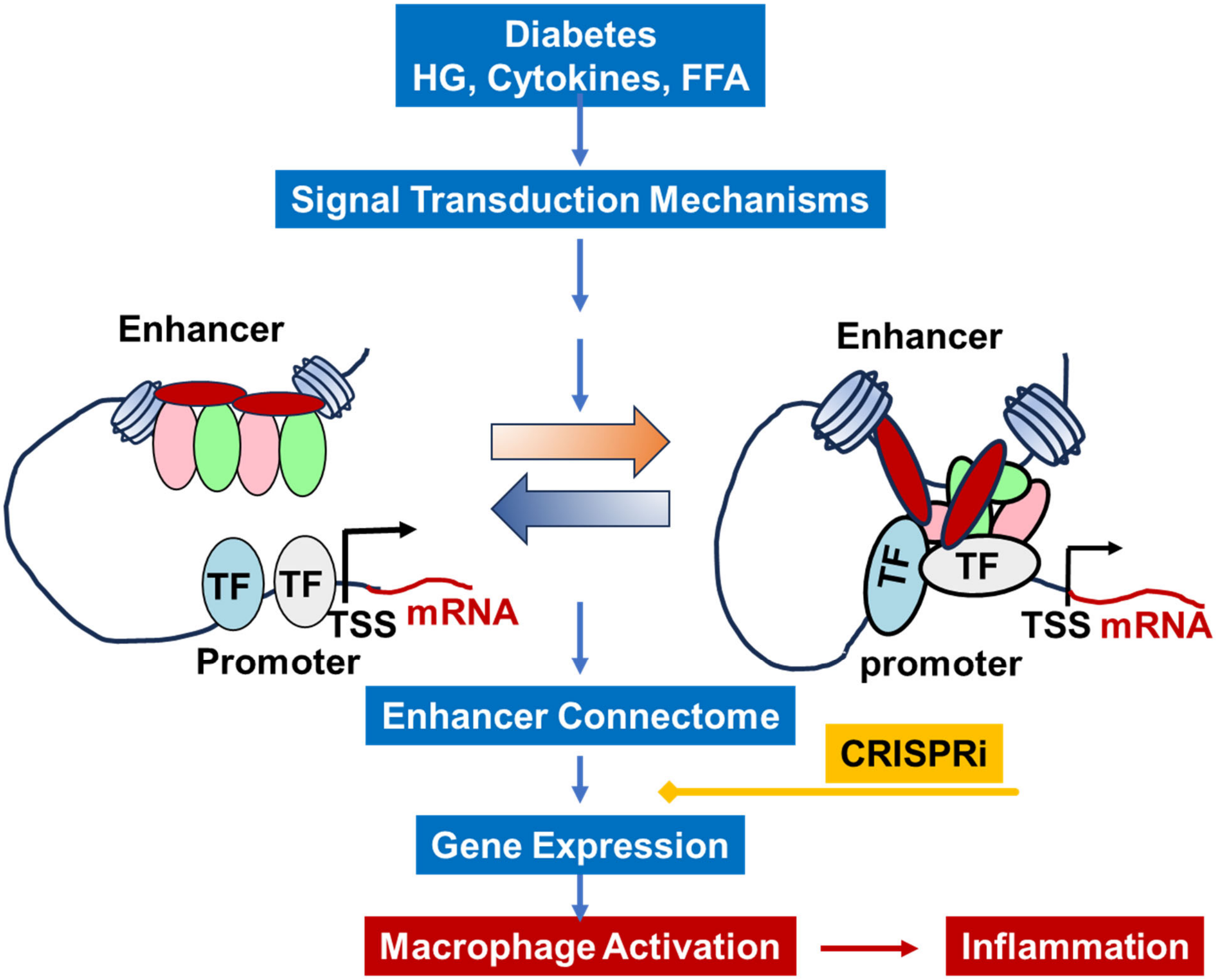


Diabetic conditions and associated pathological factors like high glucose (HG), cytokines and free fatty acids (FFA) can trigger various signal transduction mechanisms in macrophages. At the level of the nucleus and genome, this can induce extensive remodeling of the enhancer connectome, TAD boundaries and 3D chromatin architecture in macrophages, resulting in increased enhancer-bound transcription factor (TF) interactions with promoters of inflammatory (and other) genes. This can increase expression of genes that promote macrophage activation and chronic inflammation linked to diabetes complications. Targeted approaches, such as CRISPR interference (CRISPRi), can inhibit activation of key differentially regulated enhancers (such as ChemDE). Further understanding of the nuclear factors involved in such chromatin 3D remodeling under diabetic conditions can help identify new targets for epigenetic therapies for diabetes and chronic inflammatory diseases.

**Table S1**

List of TF motifs that are enriched among promoters of upregulated genes in HT and enhancers showing increased loops with gene promoters in HT

| gene | Promoter up | Enhancer loop up |
|---|---|---|
| *AHR* | Yes | No |
| *ARNT* | Yes | No |
| *IRF2* | Yes | No |
| *PPARG* | Yes | No |
| *PAX6* | Yes | No |
| *SOX9* | Yes | No |
| *REL* | Yes | Yes |
| *RELA* | Yes | Yes |
| *NR1H2* | Yes | No |
| *RXRA* | Yes | No |
| *POU5F1* | Yes | No |
| *SOX2* | Yes | No |
| *ARID3A* | Yes | No |
| *RARA* | Yes | No |
| *HIF1A* | Yes | No |
| *FOS* | Yes | Yes |
| *JUN* | Yes | Yes |
| *JUND* | Yes | Yes |
| *NR2C2* | Yes | No |
| *NRF1* | Yes | No |
| *STAT1* | Yes | No |
| *STAT2* | Yes | No |
| *ARID3B* | Yes | No |
| *LIN54* | Yes | No |
| *AR* | Yes | No |
| *ELF4* | Yes | No |
| *HNF1A* | Yes | No |
| *HSF1* | Yes | No |
| *IRF8* | Yes | No |
| *IRF9* | Yes | Yes |
| *JDP2* | Yes | Yes |
| *NEUROG2* | Yes | No |
| *NKX2-3* | Yes | No |
| *NR2F6* | Yes | No |
| *PAX4* | Yes | No |
| *SRF* | Yes | No |
| *NKX3-1* | Yes | Yes |
| *PROP1* | Yes | No |
| *RHOXF1* | Yes | No |
| *NR2F1* | Yes | No |
| *NR3C1* | Yes | No |

| | | |
|---|---|---|
| *NR3C2* | Yes | No |
| *EGR4* | Yes | No |
| *SP8* | Yes | No |
| *ZIC4* | Yes | No |
| *ZNF143* | Yes | No |
| *E2F7* | Yes | No |
| *HSF2* | Yes | No |
| *HSF4* | Yes | No |
| *IRF7* | Yes | No |
| *MEIS1* | Yes | No |
| *MEIS2* | Yes | No |
| *MYBL1* | Yes | No |
| *NFKB2* | Yes | Yes |
| *RUNX2* | Yes | No |
| *TEAD3* | Yes | No |
| *TFAP2A* | Yes | No |
| *TFAP2B* | Yes | No |
| *TFAP2C* | Yes | No |
| *OLIG1* | Yes | No |
| *CREB5* | Yes | No |
| *ALX4* | Yes | No |
| *SOX11* | Yes | No |
| *HOXD12* | Yes | No |
| *HOXC12* | Yes | No |
| *RBPJ* | Yes | No |
| *RELB* | Yes | No |
| *SIX2* | Yes | No |
| *SOX10* | Yes | No |
| *TFDP1* | Yes | No |
| *JUNB* | Yes | Yes |
| *FOSB* | Yes | Yes |
| *FOSL2* | Yes | Yes |
| *FOSL1* | Yes | Yes |
| *NR1H4* | Yes | No |
| *NR4A2* | Yes | No |
| *PPARA* | Yes | No |
| *RXRG* | Yes | No |
| *SOX15* | Yes | No |
| *IRF3* | Yes | No |
| *IRF4* | Yes | No |
| *TCF7L1* | Yes | No |
| *GLI3* | Yes | No |
| *IKZF1* | Yes | No |
| *IRF6* | Yes | No |
| *NR1I3* | Yes | No |
| *NR2C1* | Yes | No |
| *NR6A1* | Yes | No |
| *OVOL1* | Yes | No |
| *PLAGL2* | Yes | No |

| | | |
|---|---|---|
| *RFX7* | Yes | No |
| *SNAI1* | Yes | No |
| *SNAI3* | Yes | No |
| *THAP11* | Yes | No |
| *THRB* | Yes | No |
| *ZIC5* | Yes | No |
| *ZKSCAN1* | Yes | Yes |
| *ZNF136* | Yes | No |
| *ZNF140* | Yes | No |
| *ZNF460* | Yes | No |
| *ZNF75D* | Yes | No |
| *DMRT1* | Yes | No |
| *ISL1* | Yes | No |
| *PLAGL1* | Yes | No |
| *SMAD2* | Yes | Yes |
| *SMAD3* | Yes | Yes |
| *BATF* | Yes | Yes |
| *E2F3* | Yes | No |
| *EMX1* | Yes | No |
| *GLI2* | Yes | No |
| *HOXA9* | Yes | No |
| *LHX9* | Yes | No |
| *NEUROG1* | Yes | No |
| *SOX8* | Yes | No |
| *SP1* | Yes | No |
| *SP2* | Yes | No |
| *ATF2* | Yes | No |
| *BACH1* | Yes | Yes |
| *OSR2* | Yes | No |
| *TCF12* | Yes | No |
| *ZFP42* | Yes | No |
| *ZKSCAN5* | Yes | No |
| *ZNF652* | Yes | No |
| *ATF4* | Yes | No |
| *BATF3* | Yes | Yes |
| *CREB1* | Yes | No |
| *EBF1* | Yes | No |
| *GATA4* | Yes | No |
| *HOXB13* | Yes | No |
| *MAFF* | Yes | No |
| *MAFG* | Yes | No |
| *MAFK* | Yes | Yes |
| *MZF1* | Yes | Yes |
| *ONECUT1* | Yes | No |
| *PHOX2B* | Yes | No |
| *SNAI2* | Yes | No |
| *TCF3* | Yes | Yes |
| *TEAD1* | Yes | No |
| *DDIT3* | No | Yes |

| | | |
|---|---|---|
| *CEBPA* | No | Yes |
| *HAND1* | No | Yes |
| *LHX3* | No | Yes |
| *ZFX* | No | Yes |
| *GFI1B* | No | Yes |
| *MAF* | No | Yes |
| *NFE2* | No | Yes |
| *NFE2L2* | No | Yes |
| *NR2E1* | No | Yes |
| *PAX7* | No | Yes |
| *OTX1* | No | Yes |
| *NFKB1* | No | Yes |
| *PAX3* | No | Yes |
| *RFX5* | No | Yes |
| *CEBPB* | No | Yes |
| *CEBPE* | No | Yes |
| *CEBPG* | No | Yes |
| *SOX21* | No | Yes |
| *SOX1* | No | Yes |
| *BACH2* | No | Yes |
| *MAZ* | No | Yes |
| *PITX2* | No | Yes |
| *STAT5A* | No | Yes |
| *GATA5* | No | Yes |
| *PITX1* | No | Yes |
| *NFE2L1* | No | Yes |
| *OTX2* | No | Yes |
| *RFX1* | No | Yes |
| *RFX3* | No | Yes |

**Table S2**

List of TF motifs that are enriched among promoters of downregulated genes in HT and enhancers showing decreased loops with gene promoters in HT

| Gene | Promoter down | Enhancer loop down |
|---|---|---|
| *ARNT* | Yes | No |
| *DDIT3* | Yes | No |
| *CEBPA* | Yes | No |
| *MECOM* | Yes | No |
| *FOXD3* | Yes | No |
| *MAX* | Yes | No |
| *MYC* | Yes | Yes |
| *RXRA* | Yes | No |
| *VDR* | Yes | No |
| *HAND1* | Yes | No |
| *TCF3* | Yes | No |
| *NOBOX* | Yes | No |
| *EWSR1-FLI1* | Yes | No |
| *FOXH1* | Yes | No |
| *FOXO1* | Yes | No |
| *NKX2-5* | Yes | No |
| *POU2F2* | Yes | Yes |
| *SMAD2* | Yes | No |
| *SMAD3* | Yes | Yes |
| *SMAD4* | Yes | No |
| *STAT6* | Yes | No |
| *ZBTB33* | Yes | No |
| *ELK4* | Yes | No |
| *YY1* | Yes | No |
| *BACH1* | Yes | No |
| *MAFK* | Yes | No |
| *ARNTL* | Yes | Yes |
| *CREB3L2* | Yes | No |
| *DUX* | Yes | No |
| *FOXJ2* | Yes | No |
| *MLXIP* | Yes | No |
| *NPAS2* | Yes | No |
| *SIX3* | Yes | No |
| *TWIST2* | Yes | No |
| *BARHL2* | Yes | No |
| *BHLHE41* | Yes | No |
| *CENPB* | Yes | No |
| *CREB3* | Yes | No |
| *ELF5* | Yes | Yes |
| *EN1* | Yes | No |

| | | |
|---|---|---|
| *ETV6* | Yes | Yes |
| *FLI1* | Yes | No |
| *FOXI1* | Yes | No |
| *FOXL1* | Yes | No |
| *GCM1* | Yes | No |
| *GRHL1* | Yes | No |
| *GSC* | Yes | No |
| *HOXC11* | Yes | No |
| *ISX* | Yes | No |
| *KLF13* | Yes | No |
| *MLX* | Yes | No |
| *MLXIPL* | Yes | No |
| *MSX1* | Yes | No |
| *MYF6* | Yes | No |
| *NEUROD2* | Yes | No |
| *NKX2-8* | Yes | No |
| *NKX6-2* | Yes | No |
| *NR2E1* | Yes | No |
| *PAX7* | Yes | No |
| *POU4F2* | Yes | No |
| *SPDEF* | Yes | No |
| *TBX2* | Yes | No |
| *TBX21* | Yes | No |
| *TFAP4* | Yes | No |
| *TFEB* | Yes | No |
| *ZIC3* | Yes | No |
| *ZBTB18* | Yes | No |
| *MEOX2* | Yes | No |
| *MSX2* | Yes | No |
| *MSX3* | Yes | No |
| *PDX1* | Yes | No |
| *RAX* | Yes | No |
| *SHOX2* | Yes | No |
| *VAX2* | Yes | No |
| *KLF14* | Yes | No |
| *ERG* | Yes | No |
| *ETS1* | Yes | No |
| *ETV2* | Yes | Yes |
| *ETV3* | Yes | No |
| *FEV* | Yes | No |
| *GCM2* | Yes | No |
| *PAX3* | Yes | No |
| *PAX9* | Yes | No |
| *POU1F1* | Yes | No |
| *POU2F1* | Yes | No |
| *POU3F1* | Yes | No |
| *POU3F2* | Yes | No |
| *POU3F4* | Yes | Yes |
| *POU5F1B* | Yes | No |

| | | |
|---|---|---|
| *RFX2* | Yes | No |
| *RFX5* | Yes | No |
| *EOMES* | Yes | No |
| *MGA* | Yes | No |
| *TBR1* | Yes | No |
| *TBX15* | Yes | No |
| *TBX1* | Yes | No |
| *TBX4* | Yes | No |
| *TBX5* | Yes | No |
| *TFAP2B* | Yes | No |
| *BHLHE40* | Yes | No |
| *FIGLA* | Yes | No |
| *MNT* | Yes | Yes |
| *SREBF2* | Yes | No |
| *FOXC1* | Yes | No |
| *FOXC2* | Yes | No |
| *FOXO4* | Yes | No |
| *FOXJ3* | Yes | No |
| *TP73* | Yes | No |
| *GMEB2* | Yes | No |
| *MTF1* | Yes | Yes |
| *E2F1* | Yes | No |
| *TFAP2A* | Yes | No |
| *BARX1* | Yes | No |
| *DLX3* | Yes | No |
| *DLX4* | Yes | No |
| *DLX6* | Yes | No |
| *DMBX1* | Yes | No |
| *DLX2* | Yes | No |
| *EMX2* | Yes | No |
| *GBX1* | Yes | No |
| *GBX2* | Yes | No |
| *GSC2* | Yes | No |
| *HESX1* | Yes | No |
| *HOXA10* | Yes | No |
| *HOXB3* | Yes | No |
| *HOXC10* | Yes | No |
| *HOXD11* | Yes | No |
| *MYCN* | Yes | No |
| *NFYA* | Yes | No |
| *ZBTB7A* | Yes | Yes |
| *ZSCAN4* | Yes | No |
| *FOS* | Yes | No |
| *JUN* | Yes | No |
| *TFE3* | Yes | No |
| *ARNT2* | Yes | Yes |
| *ATOH1* | Yes | No |
| *ATOH7* | Yes | No |
| *BARX2* | Yes | No |

| | | |
|---|---|---|
| *CREB3L4* | Yes | No |
| *DLX5* | Yes | No |
| *DPRX* | Yes | Yes |
| *FOXN3* | Yes | No |
| *HOXA1* | Yes | No |
| *HOXA4* | Yes | No |
| *HOXA6* | Yes | No |
| *HOXA7* | Yes | No |
| *HOXB6* | Yes | No |
| *HOXB7* | Yes | No |
| *HOXB8* | Yes | No |
| *HOXC4* | Yes | No |
| *HOXC8* | Yes | No |
| *HOXD4* | Yes | No |
| *LHX1* | Yes | No |
| *LHX5* | Yes | No |
| *MAF* | Yes | Yes |
| *NFIC* | Yes | No |
| *NR5A1* | Yes | No |
| *RXRG* | Yes | No |
| *SOHLH2* | Yes | Yes |
| *SP9* | Yes | No |
| *TBX18* | Yes | No |
| *TBX3* | Yes | No |
| *TBX6* | Yes | No |
| *TLX2* | Yes | No |
| *ZBTB32* | Yes | No |
| *ZNF317* | Yes | No |
| *ZNF528* | Yes | No |
| *EBF2* | Yes | No |
| *STAT5A* | Yes | No |
| *WT1* | Yes | No |
| *ZIC1* | Yes | No |
| *ZIC2* | Yes | No |
| *ZNF281* | Yes | No |
| *BARHL1* | Yes | No |
| *BCL6* | Yes | No |
| *HES2* | Yes | No |
| *HOXD3* | Yes | No |
| *HOXD8* | Yes | No |
| *LMX1B* | Yes | No |
| *TFEC* | Yes | No |
| *BHLHE22* | Yes | No |
| *EBF3* | Yes | No |
| *MEIS1* | Yes | No |
| *MEIS2* | Yes | No |
| *NFYC* | Yes | No |
| *NKX2-2* | Yes | No |
| *PRDM4* | Yes | No |

| | | |
|---|---|---|
| *ZBTB12* | Yes | No |
| *ZNF449* | Yes | No |
| *CTCFL* | Yes | No |
| *EGR1* | Yes | No |
| *EHF* | Yes | Yes |
| *ELF1* | Yes | No |
| *ELF3* | Yes | No |
| *ETV1* | Yes | No |
| *FOXK2* | Yes | No |
| *GABPA* | Yes | No |
| *GATA1* | Yes | No |
| *GRHL2* | Yes | No |
| *MITF* | Yes | No |
| *NFYB* | Yes | No |
| *PKNOX1* | Yes | No |
| *POU2F3* | Yes | No |
| *RFX1* | Yes | No |
| *SREBF1* | Yes | Yes |
| *USF1* | Yes | Yes |
| *USF2* | Yes | Yes |
| *SOX3* | No | Yes |
| *ATF1* | No | Yes |
| *ELF4* | No | Yes |
| *MNX1* | No | Yes |
| *KLF16* | No | Yes |
| *MYBL2* | No | Yes |
| *RUNX2* | No | Yes |
| *ATF7* | No | Yes |
| *MAFB* | No | Yes |
| *GSX1* | No | Yes |
| *FOXK1* | No | Yes |
| *CDX4* | No | Yes |
| *ELF2* | No | Yes |
| *MAFA* | No | Yes |
| *NFIX* | No | Yes |
| *NKX6-3* | No | Yes |
| *ZNF75D* | No | Yes |
| *CDX1* | No | Yes |
| *GLI2* | No | Yes |
| *HES1* | No | Yes |
| *JUNB* | No | Yes |
| *NRL* | No | Yes |
| *TCFL5* | No | Yes |
| *ZNF341* | No | Yes |
| *ETV5* | No | Yes |
| *FOXA2* | No | Yes |
| *FOXP1* | No | Yes |
| *MAFF* | No | Yes |
| *MAFG* | No | Yes |

**Table S3:**

Genetic variants associated with T1D, T2D, and diabetic complications are located within HT-regulated loops, including enhancers. The genetic variants associated with different types of diabetes and their complications were retrieved from the GWAS Catalog database (https:// www.ebi.ac.uk/gwas). The HT-regulated loops with enhancers that contain any of these variants in either one of the two anchor regions were further identified and listed in the table. UAE-Urinary albumin excretion; CACAP score-Coronary artery calcified atherosclerotic plaque score.

| Category | ID | Genomic Interaction Loop | | Genotypes located within the Loop Anchor Regions | | | | | | | |
|---|---|---|---|---|---|---|---|---|---|---|---|
| | ID | anchor1 | anchor2 | DISEASE.TRAIT | SNP_ID | Chr:location | MAPPED_GENE | CONTEXT | RISK Allele Frequency | Pvalue | PMID |
| T1D only | 513 | chr6:24950000-24955000 | chr6:24950000-24955000 | T1D | 7759489 | Chr6:24950371 | RIPOR2 | intron | 0.1841 | 2.00E-06 | PMID:39749473 |
| | 1851 | chr5:35880000-35885000 | chr5:35880000-35885000 | T1D | 11954020 | Chr5:35883149 | IL7R - CAPSL | intergenic | 0.39 | 4.00E-08 | PMID:25751624 |
| | 1851 | chr5:35880000-35885000 | chr5:35880000-35885000 | T1D | 10214237 | Chr5:35883632 | IL7R - CAPSL | intergenic | 0.2752 | 2.00E-11 | PMID:39749473 |
| T1D DKD | 1612 | chr8:19065000-19070000 | chr8:19065000-19070000 | UAE in T1D | 2410601 | Chr8:19065067 | PSD3 | intergenic | 0.47 | 4.00E-06 | PMID:24595857 |
| T2D only | 139 | chr15:38560000-38565000 | chr15:38560000-38565000 | T2D | 12912777 | Chr15:38560185 | RASGRP1 | intron | 0.881128 | 5.00E-17 | PMID:39379762 |
| | 614 | chr3:23155000-23160000 | chr3:23155000-23160000 | T2D | 6780569 | Chr3:23156993 | RPL24P7 - UBE2E2-DT | intergenic | 0.83 | 4.00E-07 | PMID:23945395 |
| | 643 | chrX:153595000-153600000 | chrX:153595000-153600000 | T2D | 12010175 | ChrX:153597180 | CCNQ | intron | 0.786 | 2.00E-09 | PMID:22961080 |
| | 651 | chr16:79370000-79375000 | chr16:79370000-79375000 | T2D | 17797882 | Chr16:79373021 | RNA5SP431 - MAF | intergenic | 0.32 | 9.00E-07 | PMID:22158537 |
| | 709 | chr3:12455000-12460000 | chr3:12455000-12460000 | T2D | 4684859 | Chr3:12456902 | PPARG - TSEN2 | intergenic | 0.585781 | 1.00E-08 | PMID:39379762 |
| | 709 | chr3:12455000-12460000 | chr3:12455000-12460000 | T2D | 58691354 | Chr3:12456095 | PPARG - TSEN2 | intergenic | 0.21877 | 1.00E-09 | PMID:38374256 |
| | 1010 | chr10:86360000-86365000 | chr10:86360000-86365000 | T2D | 11201999 | Chr10:86364744 | GRID1 | intron | 0.5369 | 1.00E-08 | PMID:32541925 |
| | 1021 | chr5:119395000-119400000 | chr5:119395000-119400000 | T2D | 1105770 | Chr5:119395552 | TNFAIP8 | 3_prime_UTR | 0.42025 | 6.00E-09 | PMID:38374256 |
| | 1117 | chr9:22025000-22030000 | chr9:22025000-22030000 | T2D | 564398 | Chr9:22029548 | CDKN2B-AS1 | nc_exon | 0.56 | 1.00E-06 | PMID:17463249 |
| | 1150 | chr2:218000000-218005000 | chr2:218000000-218005000 | T2D | 76480788 | Chr2:218001763 | TNS1 | intron | 0.97431 | 1.00E-08 | PMID:38374256 |
| | 1243 | chr12:4470000-4475000 | chr12:4470000-4475000 | T2D | 75291436 | Chr12:4471074 | FGF6 - FERRY3 | intron | 0.98874 | 5.00E-08 | PMID:38374256 |
| | 1271 | chr2:145590000-145595000 | chr2:145590000-145595000 | T2D | 7609422 | Chr2:145590469 | RPL6P5 - METAP2P1 | intron | 0.4095 | 3.00E-10 | PMID:32541925 |
| | 1271 | chr2:145590000-145595000 | chr2:145590000-145595000 | T2D | 6716394 | Chr2:145593156 | RPL6P5 - METAP2P1 | intron | 0.4844 | 1.00E-13 | PMID:32541925 |
| | 1271 | chr2:145590000-145595000 | chr2:145590000-145595000 | T2D | 7609422 | Chr2:145590469 | RPL6P5 - METAP2P1 | intron | 0.37985 | 2.00E-15 | PMID:38374256 |
| | 1347 | chr3:149475000-149480000 | chr3:149475000-149480000 | T2D | 34573045 | Chr3:149478965 | TM4SF4 | intron | 0.4202 | 2.00E-11 | PMID:32541925 |
| | 1347 | chr3:149475000-149480000 | chr3:149475000-149480000 | T2D | 1802837 | Chr3:149475897 | TM4SF4 | stop_gained | 0.32625 | 1.00E-11 | PMID:38374256 |
| | 1350 | chr9:81690000-81695000 | chr9:81690000-81695000 | T2D | 2796441 | Chr9:81694033 | TLE1-DT | intron | 0.57341 | 5.00E-80 | PMID:38374256 |
| | 1403 | chr7:23470000-23475000 | chr7:23470000-23475000 | T2D | 4279506 | Chr7:23473277 | IGF2BP3 - RPS2P32 | intergenic | 0.6102 | 5.00E-08 | PMID:30297969 |
| | 1447 | chr11:43850000-43855000 | chr11:43850000-43855000 | T2D | 3736505 | Chr11:43854885 | HSD17B12 | intron | 0.30328 | 2.00E-15 | PMID:38374256 |
| | 1512 | chr3:72815000-72820000 | chr3:72815000-72820000 | T2D | 13085136 | Chr3:72816032 | SHQ1 | intron | 0.93524 | 1.00E-17 | PMID:38374256 |
| | 1547 | chr10:113030000-113035000 | chr10:113030000-11303500 | T2D | 118046020 | Chr10:113030314 | TCF7L2 | intron | 0.01105 | 2.00E-41 | PMID:38374256 |
| | 1547 | chr10:113030000-113035000 | chr10:113030000-113035000 | T2D | 72826093 | Chr10:113033836 | TCF7L2 | intron | 0.00441 | 2.00E-13 | PMID:38374256 |
| | 1547 | chr10:113030000-113035000 | chr10:113030000-113035000 | T2D | 140658764 | Chr10:113034148 | TCF7L2 | intron | 0.00772 | 3.00E-34 | PMID:38374256 |
| | 1620 | chr8:56570000-56575000 | chr8:56570000-56575000 | T2D | 62515938 | Chr8:56570454 | LINC00968 - RPL37P6 | intergenic | 0.7395 | 3.00E-08 | PMID:32541925 |
| | 1890 | chr1:219565000-219570000 | chr1:219565000-219570000 | T2D | 2820444 | Chr1:219568478 | LYPLAL1-AS1 - ZC3H11B | intergenic | 0.713 | 9.00E-21 | PMID:32541925 |
| | 1890 | chr1:219565000-219570000 | chr1:219565000-219570000 | T2D | 1415287 | Chr1:219569195 | LYPLAL1-AS1 - ZC3H11B | intergenic | 0.71817 | 1.00E-34 | PMID:38374256 |
| | 2001 | chr15:62115000-62120000 | chr15:62115000-62120000 | T2D | 8032416 | Chr15:62116871 | NPM1P47 - C2CD4B | intergenic | NR | 5.00E-09 | PMID:26818947 |
| | 1413 | chr2:217995000-218000000 | chr2:218000000-218005000 | T2D | 76480788 | Chr2:218001763 | TNS1 | intron | 0.97431 | 1.00E-08 | PMID:38374256 |
| T2D CVD | 608 | chr13:32125000-32130000 | chr13:32125000-32130000 | Atherosclerosis plaque | 543554 | Chr13:32127501 | FRY | intron | NR | 5.00E-07 | PMID:29221444 |
| Un-specified | 1350 | chr9:81690000-81695000 | chr9:81690000-81695000 | On medication | 2796441 | Chr9:81694033 | TLE1-DT | intron | 0.6197 | 6.00E-13 | PMID:39024449 |
| | 1350 | chr9:81690000-81695000 | chr9:81690000-81695000 | DM | 2796441 | Chr9:81694033 | TLE1-DT | intron | 0.631 | 5.00E-20 | PMID:39024449 |

**Table S4**

Sequences of RT-qPCR, ChIP-qPCR and 3C assay primers and details about sgRNAs used in this study.

| | **Forward** | **Reverse** |
|---|---|---|
| **RT-QPCR** | | |
| *TNF* | CCCTGAAAACAACCCTCAGA | GTCCTTTCCAGGGGAGAGAG |
| *RPLP0* | GAAACTCTGCATTCTCGCTTC | GGTGTAATCCGTCTCCACAG |
| *IL1B* | CAAAGGCGGCCAGGATATAA | CTAGGGATTGAGTCCACATTCAG |
| *IL6* | TCCTGCAGAAAAAGGCAAAG | GCCCAGTGGACAGGTTTCT |
| *CCL2* | CAAACTGAAGCTCGCACTC | CATTTCCACAATAATATCCTAG |
| *CCL11* | GGAATCCTGCACCCACTTCTT | AGGAGAATCACCAGTGGCAAA |
| *IRAK2* | GCTCACCCAATGGCTCATCT | CCTCGGCCAACACTATTCCA |
| *CCL1* | AAGCTGAAGAGAGGCAAAGAG | CAGAGCCCACAATGGAAAGA |
| *PPIB* | CAGTGGATAATTTTGTGGCCTTAG | CTCACCGTAGATGCTCTTTCG |
| **ChIP QPCR** | | |
| *CCL2* pro | TTGGAATGTGGCCTGAAGGT | AGGGTTATTTTTAAGGATTCTGCTTTC |
| *CCL*-Enh | GGCCCAGTATCTGGAATGCA | GTCAGTGCTGGCGTGAGAGA |
| ChemDE | CCCACAGCTCCACACAAA | GTTGGACAGGGATGACAGTAAA |
| *IRAK2* Enh | TGTCACAGCCCTTTCTTGTAG | CATCCCTCCAAACCTGCTC |
| **3C primers** | | |
| *ChemDE* | CATTTCCATGCCCATCCTTTATC | GAAGCCTAGCAGAACTGTAGG |
| *IRAK2* Enh | CTTCTGTTTCTAGGAAATTCAATTAGAAGG | AGACAGGAAGTTGATTGTCTATCTTTG |
| *ALDOA* | TCCTGGCAAGATAAGGAGTTGAC | ACACACGATAGCCCTAGCAGTTC |
| *IL1B* Enh | TCAAAAGCAGAAGTAGGAGGC | CACAGTTGTTGGTAAATCTTGGAG |
| *NDRG1* Enh | TGGTGTGGCAATAGGGATATTC | AAGTGTTTTCAGTGGATTTCAGTG |
| **sgRNAs** | | |
| sgPPIB | Horizon Discovery Cat# V-009250-10-002 | |
| sgNTC | Horizon Discovery Cat# U-009550-10-02 | |
| sgChemDE1 | IDT Cat # CD.Cas9.FBFM4737.AJ | |
| sgChemDE2 | IDT Cat f# CD.Cas9.ZHGR2820.AA | |
| sgChemDE3 | IDT Cat f# CD.Cas9.FBM4737.AB | |
| sgChemDE4 | IDT Cat# CD.Cas9.ZHGR2820.AB | |

IDT: Integrated DNA Technologies.
